# Live Spatial Transcriptomics Enables Longitudinal Tissue Mapping

**DOI:** 10.64898/2026.10.28.761959

**Authors:** David E. Reynolds, Sangmo Lee, Chin Nien Lee, Shailesh Senthil Kumar, Shujing Liu, Emily Huynh, Hyoyong Kim, Vladlena Hornet, Daniel Oh, Sydney Bracht, Renee-Tyler T. Morales, Phoebe Vallapureddy, Jianhua Lim, Renis J. Agosto Nieves, Junhyong Kim, Sydney Shaffer, Yoon Ho Roh, Xiaowei Xu, Yanxiang Deng, Jina Ko

## Abstract

Spatial transcriptomics has become an important technology in biomedical research by preserving the spatial context of gene expression in tissues and enabling the study of cellular states and interactions *in situ*. However, most existing methods provide only static snapshots of individual tissue specimens, requiring inference of the molecular dynamics underlying development, disease progression, and therapeutic response from different specimens rather than direct tracking in the same living tissue. Here, we introduce Live Spatial Transcriptomics (Live-ST), a technology that enables longitudinal, transcriptome-wide profiling of living tissue. Live-ST combines electroporation-based intracellular sampling and spatial barcoding to enable repeated, spatially resolved mRNA measurements in intact living tissue through next-generation sequencing. We demonstrate its capabilities by (i) benchmarking spatial mRNA capture in patterned two-dimensional cell systems, (ii) spatially profiling transcriptomes in intact, patient-derived living melanoma tissues while preserving tissue viability, and (iii) longitudinally tracking drug-response dynamics within the tumor microenvironment. Together, Live-ST enables repeated, non-destructive measurements of spatial gene expression, providing a new framework for studying dynamic tissue biology in native physiological and pathological contexts.

## Introduction

Over the past decade, spatial transcriptomics has transformed our understanding of tissue biology by enabling transcriptome-wide measurements of gene expression within intact spatial contexts.^1,2^ Unlike conventional single-cell RNA sequencing (scRNA-seq), which requires tissue dissociation, spatial transcriptomics maps cellular states and interactions directly within their native microenvironments.^3–5^ By providing insights into tissue architecture and cell-cell communication, it has been instrumental in charting embryonic development, uncovering localized disease mechanisms, and mapping the complex organization of the nervous system.^6,7^ Despite these remarkable advances, most workflows require terminal tissue processing, including fixation or freezing. Because these methods are inherently destructive, they provide only static snapshots, limiting our ability to track dynamic changes within the same tissue over time.^8^

To capture these dynamics, recent efforts have sought to integrate the temporal dimension into spatial transcriptomics. One major direction relies on physical extraction tools, such as nanostructured arrays and fluidic force microscopy designed to harvest mRNA and miRNA from living cells and fresh tissue sections.^9–14^ In parallel, metabolic RNA labeling combined with spatial transcriptomics enables spatially resolved measurements of RNA kinetics, and CRISPR-based and optogenetic technologies enable observation and control of gene expression in live, native cellular contexts.^15–18^ Other methods reconstruct temporal changes by profiling tissue collected at different time points, followed by computational integration.^19–21^ Some groups have also combined spatial transcriptomics with cellular barcoding to enable clonal tracing alongside *in situ* gene expression profiling.^22,23^ While these approaches represent significant advances and have provided valuable biological insights, they face trade-offs among molecular coverage, spatial information, and tissue viability.

Here, we present Live Spatial Transcriptomics (Live-ST), a technology that enables longitudinal, transcriptome-wide profiling of living tissue. By combining electroporation-based intracellular sampling and spatial barcoding, Live-ST enables repeated transcriptomic measurements from living tissue through next-generation sequencing (NGS) while maintaining tissue viability and preserving spatial coordinates. We first validated the platform in patterned two-dimensional (2D) cell systems, demonstrating accurate spatial mRNA capture through both optical and sequencing-based analyses. We then advanced Live-ST to patient-derived living melanoma tissues, establishing its ability to preserve tissue integrity while enabling spatial transcriptomic profiling over time. Finally, we applied our technology to monitor drug responses in patient-derived living tumor tissues, demonstrating spatiotemporal changes associated with drug treatment over time. Overall, Live-ST provides a robust framework for non-destructive spatial transcriptomics, enabling direct observation of tissue dynamics across time and space within the same living tissue.

## Results

### Live-ST workflow

Live-ST was designed to integrate electroporation-mediated intracellular RNA sampling from living tissue with deterministic spatial transcript indexing within a single workflow (Fig. 1). Our methodology includes two separate indium tin oxide (ITO) glass slides, one serving as a spatially barcoded mRNA-capture slide (anode) and the other as a tissue-supporting slide (cathode). The capture slide is brought into contact with the live tissue slice through gentle stamping, and a pulsed electric field is applied across the two slides. This extracts a fraction of intracellular molecules and electrophoretically directs RNA toward the top, spatially barcoded capture slide, building on previously demonstrated electrophoretic RNA transfer to an ITO capture surface.^24^ The sampling conditions were optimized to balance transcript recovery with preservation of cellular integrity, providing a basis for subsequent tissue culture and longitudinal spatial transcriptomic analysis. Upon completion of the transcript extraction, the intact tissue slice is immediately returned to culture for repeated sampling. The captured mRNA on the top slide undergoes reverse transcription (RT) and PCR amplification for downstream NGS. This non-destructive extraction mechanism establishes a foundation for spatiotemporal multi-sampling, enabling the tracking of transcriptomes within the same living tissue slice over time and space.

**Figure 1:**
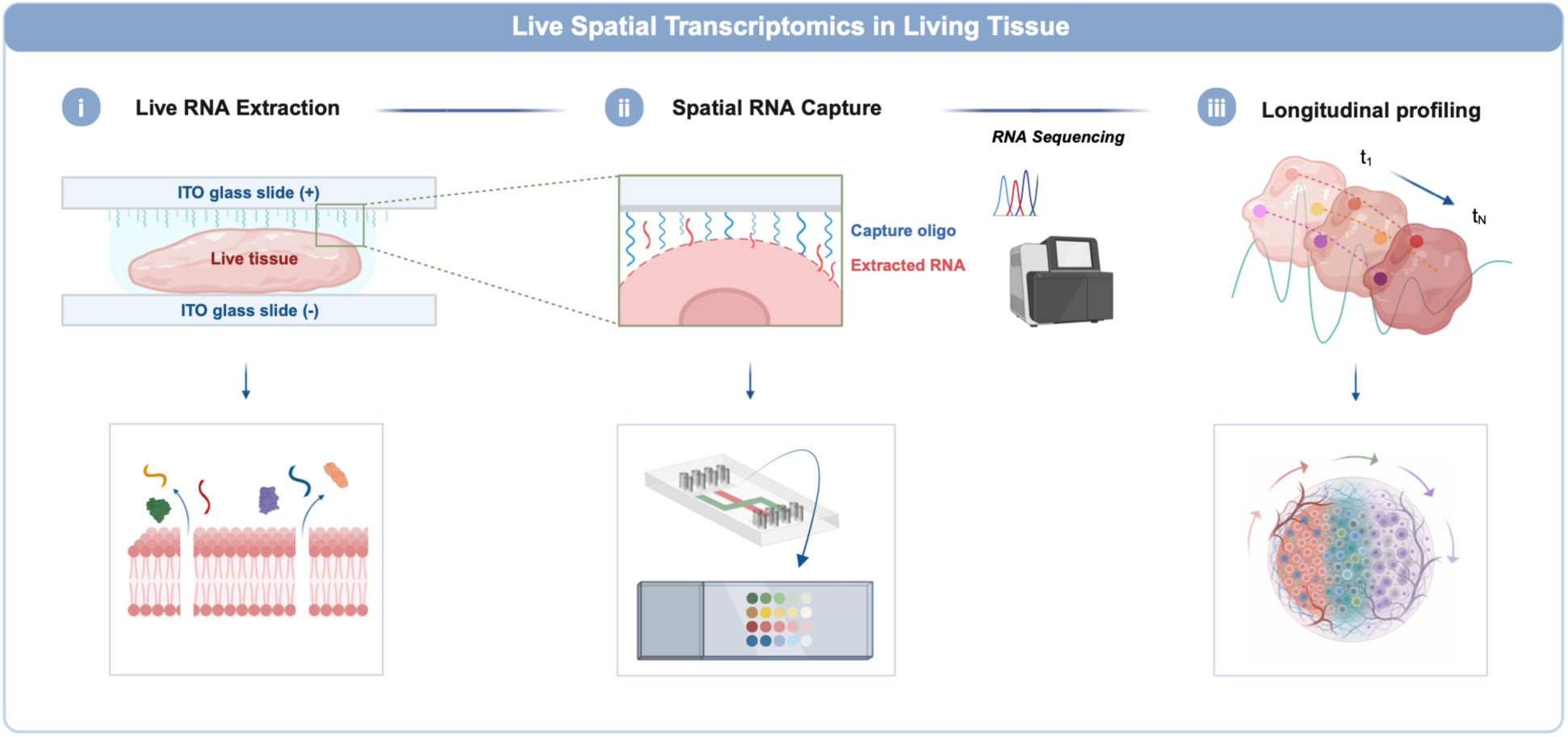
Live-ST workflow. Overview of the Live-ST workflow. A living tissue slice is placed between ITO glass slides, and a mild pulsed electric field transiently permeabilizes the cell membranes and drives negatively charged intracellular RNA toward the positively charged, spatially barcoded capture slide. Captured mRNA is processed for sequencing, and the tissue is returned to culture for repeated sampling over time. Created with BioRender.com.

### Live intracellular sampling with minimal perturbation

To optimize and parameterize intracellular sampling for patient-derived living tissue, we developed a three-dimensional (3D) melanoma-fibroblast co-culture model. Human melanoma cells (A375) and normal human dermal fibroblasts (NHDF) were embedded in a collagen I hydrogel. To replicate the dimensions of the tissue slices used in our studies, we fabricated hydrogels ∼400 µm thick using a polydimethylsiloxane (PDMS) reservoir to precisely control gel height (Fig. 2a; Supplementary Fig. 1a). For optimization, a non-barcoded ITO glass slide was placed on top of the 3D hydrogel model, and RNA extraction was performed at different electric field strengths and electroporation durations (Fig. 2b). Representative images of the model system and electrical setup are shown in Supplementary Fig. 1b. Notably, initial platform iterations utilized nanostraw arrays integrated into acrylic reservoirs; however, because lateral RNA diffusion during extraction compromised spatial resolution, we pivoted to a direct contact method in which electroporation occurs at the tissue-ITO interface to minimize transcript dispersion (Supplementary Fig. 1c–g).

**Figure 2:**
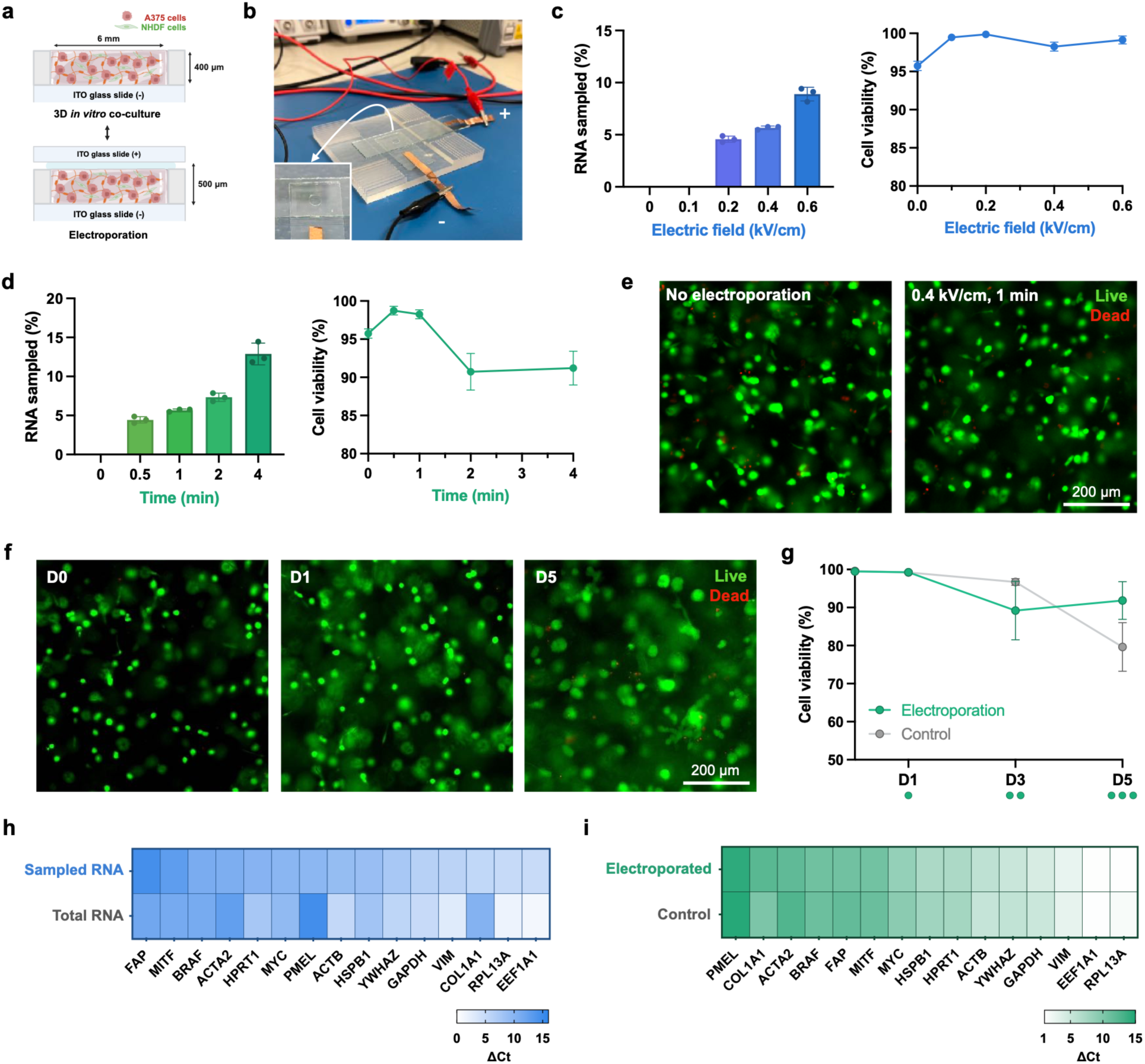
Live intracellular sampling with minimal perturbation. (**a**) Schematic of the 3D *in vitro* co-culture model. A375 and NHDF were embedded in a ∼400 µm-thick collagen I hydrogel within a 6 mm PDMS well on an ITO glass slide (−), and a second ITO glass slide (+) was placed on top for electroporation. Created with BioRender.com. (**b**) Photograph of the electroporation setup; the inset shows the hydrogel well. (**c**) RNA sampled (left) and cell viability 24 h after electroporation (right) across electric field strengths (0–0.6 kV/cm) at a fixed duration of 1 min. (**d**) RNA sampled (left) and cell viability (right) across electroporation durations (0–4 min) at 0.4 kV/cm. (**e**) Live/Dead staining 24 h after no electroporation or electroporation at 0.4 kV/cm for 1 min. Live cells are shown in green and dead cells in red. (**f**) Live/Dead staining during repeated sampling at D0, D1, and D5. (**g**) Cell viability of electroporated samples and non-electroporated controls at baseline (D0) and after one (D1), two (D3), and three (D5) sampling events; dots below the x-axis indicate the cumulative number of sampling events. (**h**) Relative expression (ΔCt) of target transcripts in RNA sampled by Live-ST and in total RNA from whole-cell lysis. (**i**) ΔCt of the same transcripts in electroporated cells lysed 48 h after sampling and in non-electroporated controls. Data in (**c**,**d**,**g**) are shown as mean ± SD (*n* = 3 technical replicates). *P* values in (**g**,**i**) were calculated with unpaired two-tailed *t*-tests. Scale bars: 200 µm (**e**,**f**).

We optimized the electroporation parameters on our 3D hydrogel model by evaluating RNA extraction efficiency alongside cell viability. With a constant 1 min duration, increasing the electric field enhanced RNA extraction efficiency from 0% to ∼9% while maintaining high post-electroporation viability (>95% after 24 h; Fig. 2c). We selected a pulsed electric field of 0.4 kV/cm (50 Hz, 200 µs pulse width, 1.0% duty cycle), which yielded ∼6% RNA extraction per cell, an amount sufficient for sequencing without compromising cell health.^9,25^ Crucially, the low-duty-cycle micro-pulsing reduces cumulative electrical exposure, potentially limiting heating and electrochemical effects. At 0.4 kV/cm, extending electroporation to 2–4 min increased RNA extraction yield from ∼6% to ∼13%, but viability declined to ∼91% (Fig. 2d). We therefore used a 1-min duration at 0.4 kV/cm in all subsequent live hydrogel and cell-line experiments; cell viability imaging 24 h post-treatment confirmed no detectable morphological or viability differences compared to non-electroporated controls (Fig. 2e). Compared to our optimized ITO direct-contact approach, nanostraw-based sampling yielded lower overall RNA extraction efficiency (Supplementary Fig. 1h). This difference is largely because nanostraw parameters were implemented from literature protocols rather than specifically optimized for our tissue model.^25^

To determine whether repeated sampling alters long-term cell health, we assessed cell viability at baseline (D0) before electroporation, and immediately after repeated sampling of the same samples on days 1, 3, and 5 (D1, D3, and D5). Cells subjected to up to three consecutive sampling rounds maintained high viability (Fig. 2f). Quantification showed no statistically significant difference in cell viability between electroporated samples and non-electroporated controls over the 5-day course (Fig. 2g; unpaired t-test, *P* = 0.837), confirming that our platform supports repeated, non-destructive sampling over extended culture periods. Next, we evaluated the relative representation of selected transcripts comprising lineage markers, stress-response, and housekeeping genes in sampled RNA obtained using our extraction method compared with total RNA from whole-cell lysis by quantitative PCR (primers in Supplementary Table 1).^26–31^ The relative expression levels (ΔCt) obtained via non-destructive extraction closely mirrored those from total RNA, showing a positive correlation (Pearson’s *r* = 0.73; Fig. 2h). Furthermore, to confirm that sampling does not perturb baseline gene expression, we lysed the electroporated cells 48 h post-extraction and compared their gene expression with that of non-electroporated controls. The post-sampling expression profiles (ΔCt) showed no statistically significant differences from non-electroporated controls (unpaired t-test, *P* = 0.851), suggesting minimal perturbation of the tested transcripts following intracellular sampling (**Fig. 2i**). Collectively, these optimization steps establish a robust framework for live intracellular sampling in a 3D hydrogel model, balancing efficient transcript yield with long-term cellular preservation.

### Spatial preservation of captured RNA

To evaluate our technology’s ability to preserve mRNA spatial localization after extraction, we conducted a series of studies using both optical and sequencing-based readouts. We first assessed the platform using patterned, fixed A549 cells with an optical readout before progressing to tissue samples. For these experiments, we employed an ITO glass slide functionalized with amine-modified poly(T) capture probes to directly visualize mRNA capture. Following mRNA capture and RT, the glass slide was treated to remove unextended capture probes, leaving only the RNA-cDNA hybrids. These hybrids were then stained with SYBR Gold and visualized by fluorescence microscopy. We validated the optical workflow by incubating purified A549 total RNA on the capture surface. Fluorescence intensity increased with RNA input, with 1,000 ng of RNA producing a ∼9-fold higher signal than the no-RNA control, demonstrating assay sensitivity (Extended Data Fig. 1a).

For the cell-based study, live A549 cells were seeded into laser-cut acrylic wells patterned in the shape of a “T” and embedded within a thin PDMS reservoir on an ITO glass slide. After 24 h, the cells were fixed and the acrylic mold was removed, leaving only the patterned cells on the glass slide (Extended Data Fig. 1b). Comparison of stamped and unstamped samples showed no noticeable distortion of the cellular pattern, indicating that the direct contact process did not visibly alter the spatial arrangement of cells (Extended Data Fig. 1c). To assess spatial preservation of mRNA, an ITO glass slide functionalized with amine-modified poly(T) capture probes was placed over the fixed cells using the same direct-contact and electroporation setup as for live intracellular sampling, and briefly electroporated to extract and capture intracellular mRNA. Following RT and staining, a distinct “T”-shaped pattern was observed that closely matched the original cell geometry. In contrast, non-electroporated control samples produced only a faint, unrecognizable signal, confirming that electroporation is essential for efficient mRNA extraction and capture (Extended Data Fig. 1d). Overlaying the RNA-cDNA hybrid and fixed-cell images confirmed strong spatial fidelity (Extended Data Fig. 1e). To quantify this, binary masks were generated using intensity thresholding and morphological cleaning. The captured cDNA reproduced the T-shaped pattern of the cells (Dice coefficient = 0.88), with 96% of the cDNA signal area lying within the cell-covered area (Extended Data Fig. 1f). At the cell boundary, the cDNA signal fell to half its intensity within ∼19 µm (≤32 µm across reference-window choices), less than both the 50 µm barcode spots and the 50 µm interspot spacing of the spatial barcoding array used in subsequent experiments, indicating that lateral RNA diffusion during capture is minimal at the array’s resolution. Together, these results confirm that mRNA spatial localization is preserved during extraction and transfer to the capture surface.

### Spatial transcriptomic profiling of living tissue

We next adapted Live-ST to living tissue using patient-derived melanoma samples (patient characteristics in Supplementary Table 2). Biopsies were taken using a 5 mm punch and sectioned into 400 µm slices with a vibratome (Supplementary Fig. 2a,b). To ensure robust intracellular extraction despite variability in tissue geometry and thickness over culture time, we engineered an improved mechanical setup. A manual z-axis dovetail stage was fitted with custom 3D-printed ITO glass slide holders, and a load cell was positioned directly beneath the tissue-supporting slide to provide real-time feedback and precisely detect tissue contact (Supplementary Fig. 2c,d). Before confirming spatial mRNA capture in live tissue, we evaluated tissue health 24 h post-electroporation (Fig. 3a). Live/Dead fluorescence imaging revealed comparable high cell viability between unsampled (Live-ST (−)) and sampled (Live-ST (+)) slices (Fig. 3b). Metabolic activity quantified via MTS assay showed comparable absorbance between the two conditions (Fig. 3c).

**Figure 3:**
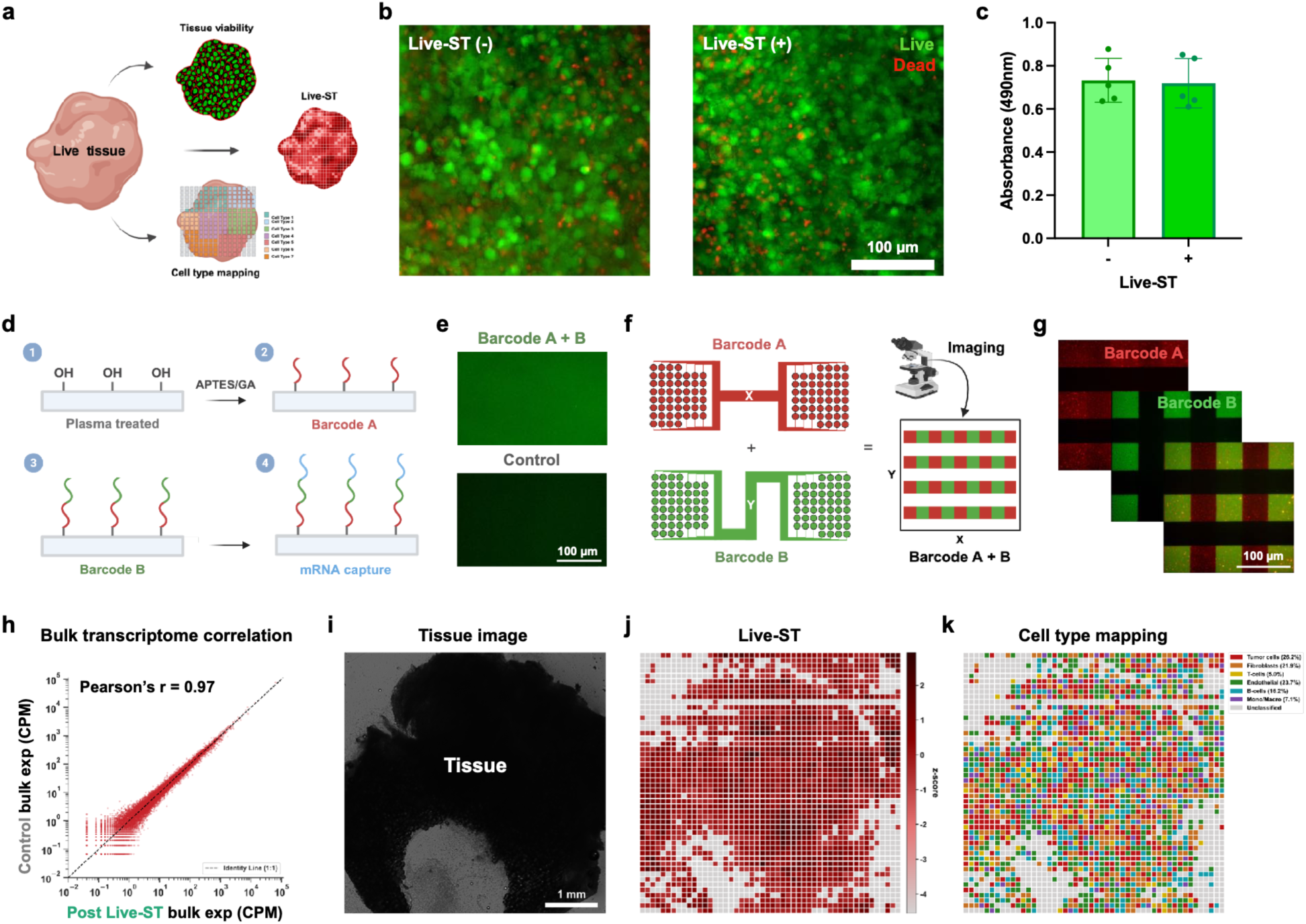
Spatial transcriptomic profiling of living tissue. (**a**) Schematic of tissue-level analyses: tissue viability, Live-ST spatial profiling, and cell type mapping. Created with BioRender.com. (**b**) Live/Dead staining of melanoma slices (Patient 1) 24 h after Live-ST (+) sampling or without sampling (Live-ST (−)). Live cells are shown in green and dead cells in red. (**c**) Metabolic activity of Live-ST (−) and Live-ST (+) slices measured by MTS assay (CellTiter 96 AQueous One Solution Cell Proliferation Assay, Promega; absorbance at 490 nm). (**d**) Surface functionalization of the capture slide: plasma treatment, 3-aminopropyltriethoxysilane (APTES)/glutaraldehyde-mediated covalent immobilization of amine-modified barcode A, enzymatic ligation of barcode B containing a UMI and poly(T) sequence, and mRNA capture. (**e**) Fluorescence of ligated barcode B on barcode A-functionalized (Barcode A + B) and control slides. (**f**) Combinatorial spatial barcoding. Barcodes A and B are delivered through orthogonal sets of 50 parallel PDMS microfluidic channels along the x- and y-axes, generating a 50 × 50 grid of 2,500 uniquely barcoded spots. (**g**) Fluorescence image of crossed barcode A (red) and barcode B (green) channels, showing the checkerboard pattern. (**h**) Bulk expression (counts per million; CPM) of Live-ST-sampled versus control tissue (Patient 1) 48 h after sampling. The dashed line indicates the identity line. (**i**) Bright-field image of the melanoma slice from Patient 2; a 1 mm biopsy punch at one side served as a positional landmark. (**j**) Live-ST spatial map of UMI counts per spot (z-score of log(UMI + 1)); grey spots had fewer than 25 uniquely mapped UMIs and failed quality control. (**k**) Spatial map of the dominant cell type in each spot, inferred by cell type mapping with a scRNA-seq reference. Percentages indicate the proportion of tissue spots assigned to each cell type; grey spots were unclassified. Data in (**c**) are shown as mean ± SD (*n* = 5 slices). Scale bars: 100 µm (**b**,**e**,**g**), 1 mm (**i**).

For sequencing-based spatial readout, the capture slide was prepared by 3-aminopropyltriethoxysilane (APTES) silanization and glutaraldehyde crosslinking, enabling covalent immobilization of amine-modified oligonucleotides (barcode A) onto the surface. The 5’-phosphorylated oligonucleotides (barcode B), containing a unique molecular identifier (UMI) and a poly(T) sequence, were enzymatically ligated to barcode A (Fig. 3d). Successful surface functionalization was confirmed by the selective fluorescence signal of barcode B, which was observed only on barcode A-functionalized slides following *in situ* ligation (Fig. 3e). To achieve deterministic spatial barcoding, we adopted a combinatorial indexing strategy utilizing PDMS microfluidic devices and barcode oligonucleotide sequences, following approaches established in previous studies (Fig. 3f).^4,32,33^ By sequentially applying two sets of 50 parallel channels along the x- and y-axes, we generated a precisely aligned 50 × 50 grid comprising 2,500 unique spatial spots, which was visually verified by a distinct checkerboard fluorescence pattern (Fig. 3g).

Bulk sequencing of tissue slices 48 h after Live-ST further confirmed minimal transcriptional disruption, showing a strong correlation between sampled and control tissues (Pearson’s *r* = 0.97; Fig. 3h). These findings support minimal transcriptional perturbation following Live-ST sampling. Before applying the barcoded slide to tissue, we evaluated its sequencing-based spatial readout in patterned 2D cells. Because the barcoded region of the capture slide covers a 5 × 5 mm area, we patterned live A549 cells within four 1 mm PDMS reservoirs positioned inside this region on an ITO glass slide. After cell seeding and culture, the PDMS reservoirs were removed, leaving only the patterned A549 cells on the glass slide surface (Extended Data Fig. 2a). A reference spatial map was generated by reconstructing the expected cell distribution from the initial bright-field image and served as the ground truth for spatial comparison (Extended Data Fig. 2b). The Live-ST sequencing output displayed spatial distributions closely aligned with the reference map (Extended Data Fig. 2c). Quantitative intensity line profiles across four cross-sectional barcode coordinates (X = 15, 35; Y = 15, 35) confirmed matching peak locations and similar slopes between reference and experimental data (Extended Data Fig. 2d). Read depth, uniquely mapped reads, and per-spot UMI and gene counts are shown in Extended Data Fig. 2e–g. Cell viability assessed 24 h after sampling remained high (>95%) in both sampled and unsampled cells (Extended Data Fig. 2h,i).

To confirm that Live-ST preserves spatial mRNA patterns in living tissue, we created a positional landmark by taking a 1 mm biopsy punch from one side of a melanoma sample from Patient 2 (Fig. 3i). After applying Live-ST to the sample, we reconstructed the sequencing data and confirmed that both the raw UMI counts and normalized expression maps accurately recapitulated the physical contours and structural landmark of the tissue (Fig. 3j; Supplementary Fig. 3a,b). Sequencing quality metrics for this tissue are shown in Supplementary Fig. 3c–e. We then mapped the spatial spots to specific cell types using an scRNA-seq reference dataset.^34^ Cell type mapping revealed a heterogeneous tumor-immune microenvironment composed of tumor cells (26.2%), endothelial cells (23.7%), fibroblasts (21.9%), B cells (16.2%), monocytes/macrophages (7.1%), and T cells (5.0%). Consistent with the 50 µm spot size spanning multiple cells, most spots contained inferred contributions from several cell types (Fig. 3k; Supplementary Fig. 3f). To assess whether the inferred composition was representative of melanoma tissue, we compared it with flow cytometry of dissociated cells from a different patient, with proportions scaled to sum to 100% across the six cell types (Supplementary Fig. 3g). Cells were stained with markers for tumor cells (SOX10, HMB45), fibroblasts (α-SMA), endothelial cells (CD31), and immune populations (CD45, CD3, CD19, and CD11b) (Supplementary Fig. 3h–k). Live-ST detected all six major populations identified by flow cytometry, with tumor cells the most abundant in both, although it underrepresented tumor cells (26.2% vs. 46.1%) and overrepresented endothelial cells (23.7% vs. 4.9%) and B cells (16.2% vs. 6.6%). These differences may reflect the different measurement scales of Live-ST (dominant cell type per spatial spot) and flow cytometry (individual cell proportions), as well as inter-patient variation, cell type-specific differences in transcript abundance and extraction, tissue dissociation biases, or incomplete representation in the scRNA-seq reference. In addition, flow cytometry categories were defined by single markers and only approximate the transcriptome-defined Live-ST categories; for example, CD11b also marks neutrophils and other myeloid cells, and α-SMA labels activated fibroblasts but not all fibroblasts. Together, these results demonstrate that Live-ST recapitulates tissue geometry and identifies the major cell populations when applied to living tissue.

### Longitudinal spatial profiling of drug responses in living tissue

To demonstrate our platform’s ability to track tissue changes over time, we profiled drug responses in patient-derived melanoma samples. We used *BRAF* V600E-mutant melanoma tissue from a single patient (Supplementary Table 2), cored with a 6 mm biopsy punch and sectioned into 400 µm slices. Two slices were treated with the MEK inhibitor trametinib (10 nM) and the BRAF inhibitor dabrafenib (100 nM) (Treated 1 and 2) and two slices were maintained without drug (Nontreated 1 and 2).^35^ Each slice was profiled with Live-ST one day before treatment (D-1) and again on days 1 and 3 after treatment (D1 and D3), yielding 12 spatial datasets from four living tissue slices (Fig. 4a; Supplementary Fig. 4a). Concurrently with the sequencing study, additional drug-treated and nontreated slices from the same patient were cultured with and without Live-ST sampling to serve as viability controls. To align data across each time point, we created a positional landmark with a 1 mm biopsy punch at the tissue edge, which allowed each slice to be registered to the 5 × 5 mm barcoded area at every sampling event (Fig. 4a,b). Hematoxylin and eosin (H&E) staining of an adjacent tissue slice confirmed densely cellular melanoma tissue (Fig. 4c).

**Figure 4:**
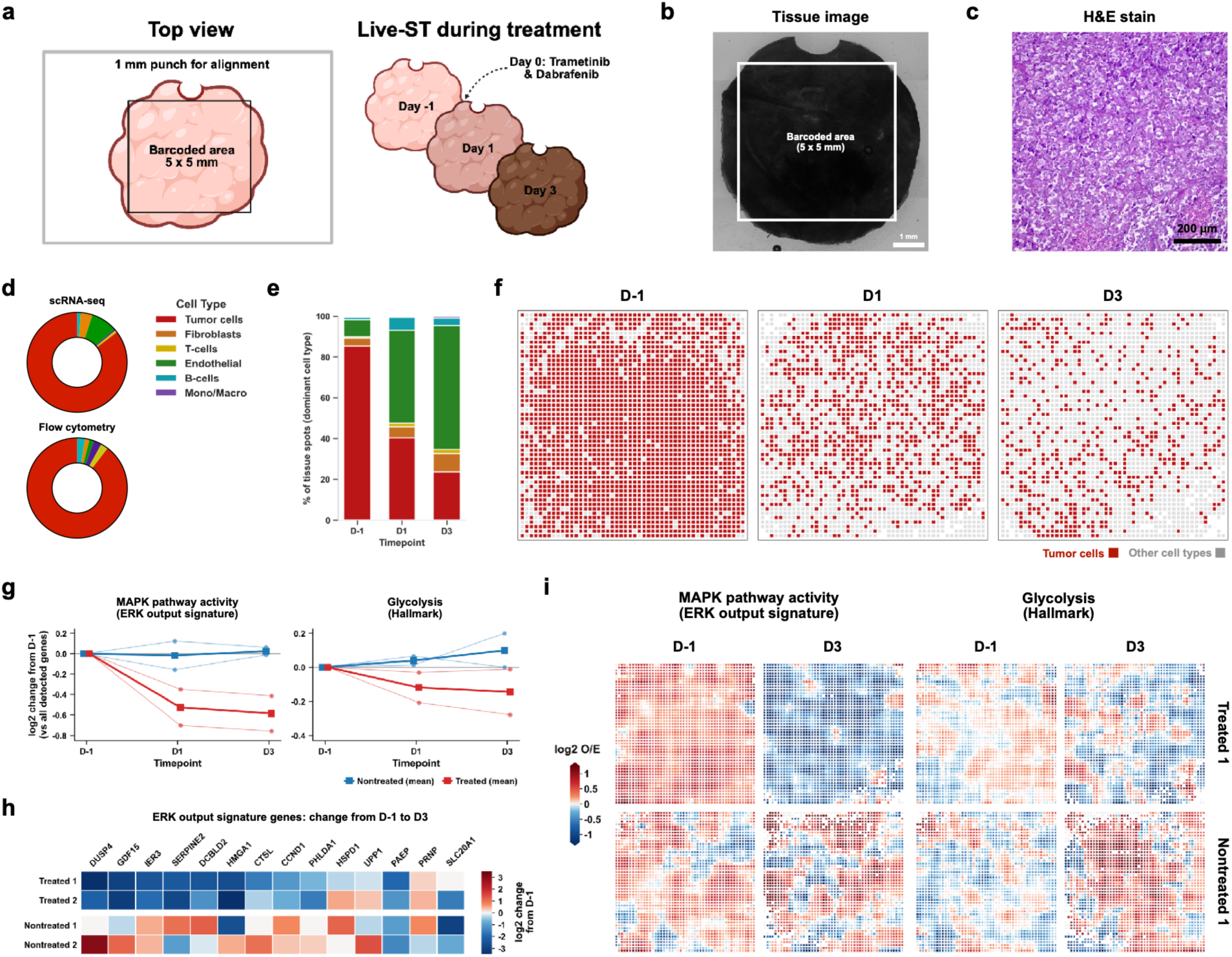
Longitudinal spatial profiling of drug responses in living tissue. (**a**) Experimental design. A 1 mm biopsy punch at the tissue edge served as a landmark for aligning each slice to the 5 × 5 mm barcoded area at every sampling event (left). Slices were profiled by Live-ST at D-1, D1, and D3, with trametinib (10 nM) and dabrafenib (100 nM) added at D0 (Treated 1 and 2) or omitted (Nontreated 1 and 2) (right). Created with BioRender.com. (**b**) Bright-field image of a 6 mm melanoma slice from Patient 4 with the 5 × 5 mm barcoded area outlined. (**c**) H&E staining of the melanoma tissue. (**d**) Cell type composition of Treated 1 at baseline (D-1), estimated from Live-ST spots mapped with the scRNA-seq reference (top, “scRNA-seq”) and measured by flow cytometry of the same patient sample (bottom). Flow cytometry proportions were scaled to sum to 100% across the six cell types. (**e**) Proportion of tissue spots by dominant cell type in Treated 1 at D-1, D1, and D3. (**f**) Spatial maps of tumor-dominant spots (red) and spots dominated by other cell types (grey) in Treated 1 at D-1, D1, and D3. (**g**) Change from D-1 in MAPK pathway activity (ERK output signature; left) and glycolysis (Hallmark; right) module scores, expressed as log_2_ change relative to all detected genes. Thin lines show individual tissues; thick lines show the mean of treated (red) or nontreated (blue) tissues. (**h**) Log_2_ change in expression from D-1 to D3 in each tissue for the 14 ERK output signature genes with at least 5 UMIs at D-1 and D3 combined in every tissue (selected by depth, not by change), ordered by the treated minus nontreated difference. (**i**) Spatial maps of MAPK pathway activity and glycolysis (log_2_ observed/expected) at D-1 and D3 in Treated 1 and Nontreated 1. In (**g**,**h**), each tissue’s three samples were downsampled to the same depth before computing changes from D-1, and (**h**) shows each gene’s own change. Maps in (**i**) compare each 7 × 7-spot window with the count expected from its depth (Methods). Scale bars: 1 mm (**b**), 200 µm (**c**).

These viability controls showed that Live-ST sampling itself did not compromise tissue viability over the course of the study. As expected, metabolic activity in the drug-treated controls declined modestly relative to nontreated controls (fold change of ∼0.87 at D1 and ∼0.82 at D3), consistent with an on-target drug effect (Supplementary Fig. 4b).^36^ Live/Dead imaging at D-1 and after the final time point (D3) showed comparable viability between Live-ST (+) and Live-ST (−) slices (Supplementary Fig. 4c,d). Sequencing metrics and per-spot UMI and gene counts for all 12 libraries are shown in Supplementary Fig. 5a,b, and UMI maps of Treated 1 at D-1 in Extended Data Fig. 3a,b. Pseudobulk gene expression showed moderate positive correlations between time points within each tissue (Pearson’s *r* = 0.50–0.67; Supplementary Fig. 5c,d).

We next examined how cellular composition changed over time within the same tissue. At baseline (D-1), Live-ST estimated that 85.3% of spots in Treated 1 were dominated by tumor cells, closely matching the 89.6% tumor fraction measured by flow cytometry on the same patient sample (Fig. 4d). Following treatment, the fraction of tumor-dominant spots in Treated 1 fell from 85.3% (D-1) to 40.3% (D1) and 23.5% (D3), while the fraction of endothelial-dominant spots rose from 8.4% to 45.5% and 60.7%, respectively (Fig. 4e,f; Extended Data Fig. 3c). Because these measurements were taken from the same living tissue, these shifts reflect changes within a single sample rather than differences between adjacent sections. However, tumor-dominant spots also decreased in nontreated slices, so these compositional changes cannot be attributed to drug treatment alone. We therefore focused subsequent analyses on pathway activity.

We asked whether Live-ST could resolve the expected pharmacodynamic response to treatment. Mitogen-activated protein kinase (MAPK) pathway activity, measured using an extracellular signal-regulated kinase (ERK) output signature, decreased in both treated tissues by D1 (mean log_2_ change from D-1 of approximately −0.53) and remained suppressed at D3 (approximately −0.59), whereas nontreated tissues remained near baseline (approximately −0.02 at D1 and +0.02 at D3). This reduction in drug-treated samples is consistent with the expected inhibition of MAPK signaling by combined BRAF and MEK inhibition. The Hallmark glycolysis signature decreased in treated tissues (approximately −0.12 at D1 and −0.15 at D3) and slightly increased in nontreated tissues (approximately +0.04 and +0.10), which is consistent with reduced glycolytic gene expression downstream of MAPK inhibition (Fig. 4g).^37,38^ At the gene level, canonical ERK targets including *DUSP4*, *GDF15*, *IER3*, *DCBLD2*, and *CCND1* were consistently down-regulated from D-1 to D3 in both treated tissues, but not in nontreated tissues (Fig. 4h). Spatial maps further revealed that ERK-target expression was suppressed across nearly the entire treated tissue by D3, whereas the nontreated tissue retained regionally heterogeneous MAPK activity. The glycolysis signature showed a modest, regionally variable increase in the nontreated tissue by D3 but not in the treated tissue (Fig. 4i). We next examined tumor necrosis factor-α (TNF-α) signaling via nuclear factor-κB (NF-κB), an inflammatory pathway implicated in melanoma resistance to MAPK inhibitors.^39,40^ To distinguish NF-κB-associated transcriptional changes from ERK output, we excluded genes shared between the two signatures. Treatment also reduced this signature in treated tissues (mean log_2_ change of approximately −0.42 at D1 and −0.32 at D3) compared with smaller changes in nontreated tissues (approximately −0.09 and −0.10), suggesting dampened inflammatory signaling within the tumor tissue (Extended Data Fig. 3d,f). The reduction was smaller at D3 than at D1, consistent with partial recovery of NF-κB-associated transcription under sustained MAPK inhibition.^41^

Because tumor composition changed over time, we tested whether these pathway changes simply reflected the loss of tumor-dominant spots. When restricted to tumor-dominant spots, module score changes closely tracked those computed across all tissue spots, falling near the 1:1 line for the ERK output, glycolysis, and TNF-α/NF-κB signatures (Extended Data Fig. 3e). Thus, the decrease in MAPK activity reflects transcriptional changes within tumor regions rather than shifts in cellular composition alone. Analysis of the most abundant transcripts and their associated Gene Ontology terms showed shared immune and stress-response programs across conditions, with host defense response and mRNA stability terms enriched only in treated tissue at D3 (Extended Data Fig. 3g–j). Together, these results demonstrate that Live-ST can longitudinally track drug responses in the same living tissue, capturing spatially resolved changes in pathway-associated gene expression within the tumor-immune microenvironment that cannot be directly tracked in the same specimen using conventional endpoint spatial assays.

## Discussion

The principal advance of Live-ST is the ability to follow spatial gene expression within the same living tissue over time, rather than reconstructing temporal changes from independently collected specimens. This distinction is particularly important in heterogeneous tissues, where differences in cellular composition and local microenvironment can confound comparisons between samples. Live-ST pairs electroporation-mediated intracellular RNA sampling with spatial barcoding to enable repeated, transcriptome-wide profiling in living tissue. By utilizing an optimized dual-substrate ITO direct-contact setup, Live-ST restricts transcript diffusion during sampling while preserving cell viability. Crucially, parameter tuning allows controlled mRNA extraction, enabling multi-day longitudinal sampling without perturbing baseline cellular states or transcriptomic profiles.

Applied to *ex vivo*-treated *BRAF*-mutant melanoma slices, Live-ST detected rapid, sustained suppression of MAPK/ERK output and reduced glycolytic and NF-κB inflammatory signaling. Importantly, these pathway changes persisted within tumor regions independent of compositional shifts. By repeatedly profiling the same tissue, Live-ST allows drug-associated transcriptional changes to be followed relative to each specimen’s own baseline, revealing their temporal persistence and spatial distribution. Ultimately, Live-ST advances spatial transcriptomics from a static endpoint assay into a dynamic, longitudinal platform, enabling the tracking of spatiotemporal cellular interactions and drug response dynamics within intact living microenvironments.

Although Live-ST enables longitudinal profiling in living tissue, several limitations remain. First, only a small fraction of intracellular RNA is extracted at each sampling event, which restricts the detection of low-abundance transcripts. However, coupling spatial barcoding with nanowires or nanoneedles could boost RNA yield without sacrificing spatial resolution. Second, per-spot transcript capture decreased at later sampling time points. Improving the consistency of capture across repeated sampling events will strengthen quantitative longitudinal analyses. Third, the drug-response study used tissue from a single patient who was receiving encorafenib and binimetinib at the time of resection, so prior BRAF and MEK inhibitor exposure may have shaped the *ex vivo* response. Studies in additional patients, including treatment-naive tissue, will be needed to generalize these findings. Additionally, maintaining *ex vivo* tissue viability is challenging, particularly because sampling requires removing tissue from culture. Developing integrated, scalable culture platforms would allow Live-ST to be performed directly within the culture system, minimizing handling-related perturbations. Live-ST also currently lacks single-cell resolution, which future iterations could achieve by using higher-density (sub-10 µm) barcode arrays combined with image-based cell segmentation. Despite these constraints, Live-ST establishes a foundation for tracking continuous transcriptomic dynamics over time. This approach offers future potential for deciphering developmental biology and organogenesis, tracking disease progression in the same tissue, and guiding therapeutic discovery in intact biological systems.

## Methods

### Cell culture

A375 cells (ATCC, CRL-1619), A549 cells (ATCC, CRM-CCL-185), and NHDF (ATCC, PCS-201-012) were cultured in DMEM supplemented with 10% FBS and 1% penicillin-streptomycin. All cells were maintained at 37 °C with 5% CO2.

### Tissue preparation and culture

Fresh melanoma tissues were obtained from the Hospital of the University of Pennsylvania, Department of Pathology and Laboratory Medicine. All procedures were approved by the Institutional Review Board at the University of Pennsylvania (protocol no. 703001). Patient characteristics are listed in Supplementary Table 2. The fresh tissues were maintained in RPMI-1640 medium until processing. Tumors were washed three times with DPBS without calcium and magnesium, containing 2% penicillin-streptomycin, and cored with a 5 mm (spatial mapping) or 6 mm (longitudinal drug-response profiling) biopsy punch (Miltex). Cores were embedded in 1.5% (w/v) agarose in molecular-grade water and sectioned into 400 µm slices on a vibratome (Precisionary Instruments, VF-310-0Z; blade speed 10, oscillation 10). Slices were collected in DPBS containing 2% penicillin-streptomycin, washed three times in DPBS without calcium and magnesium containing 2% penicillin-streptomycin, and cultured on Transwell inserts (0.4 µm pore; Fisher Scientific, 08-771) in 6-well plates in RPMI 1640 supplemented with 10% FBS, 1% penicillin-streptomycin, and 100 IU/mL recombinant human IL-2 (PeproTech, AF-200-02) at 37 °C with 5% CO2. Medium was replaced daily.

### 3D hydrogel fabrication

PDMS(Sylgard 184, Corning) was mixed at a 10:1 base-to-curing-agent ratio, spin-coated onto glass slides at 500 rpm for 10 s, and cured at 50 °C for 1 h, producing ∼400 µm-thick layers as measured with a caliper. Reservoirs were punched with a 6 mm biopsy punch, and the PDMS layer was placed onto ITO-coated glass slides (1.1 mm thick, 30–60 Ω/sq; MSE, GL0346) that had been cleaned sequentially with acetone, isopropanol, and deionized (DI) water. A375 cells and NHDF were mixed at a 1.5:1 ratio in DMEM containing 10% FBS and 1% penicillin-streptomycin and combined with collagen I (Corning, CB40236; final concentration 1.5 mg/mL), serum-free DMEM, and 1 N NaOH (final concentration 0.01 M) for neutralization. Each reservoir was filled with the cell-collagen mixture (∼10,000 cells per well) until level with the top of the PDMS layer and allowed to gel at 37 °C for 30 min. DMEM supplemented with 10% FBS and 1% penicillin-streptomycin was then added, and hydrogels were cultured as described above.

### Electroporation setup

Two ITO-coated glass slides served as electrodes, with the sample-supporting slide as the cathode and the top slide as the anode. Electrical contact was made with copper tape (McMaster-Carr). Square-wave pulses were produced using a DDS signal generator (Koolertron) and a programmable DC power supply (RIGOL Technologies, DP711), and waveforms were monitored with a digital oscilloscope (RIGOL Technologies, DHO814). Electric field strength was calculated as the applied voltage divided by the electrode gap.

For tissue experiments, the capture slide was mounted in custom 3D-printed holders (University of Pennsylvania Libraries’ Holman Biotech Commons) on a manual z-axis dovetail stage (Amazon, B0G8D6V525). A load cell (DigiKey, 1568-14727-ND) positioned beneath the tissue-supporting slide was read through an HX711 amplifier (DigiKey, 1568-13879-ND) and an Arduino UNO R3 to detect contact between the capture slide and the tissue as the stage was lowered.

### Intracellular sampling optimization

After 24 h of culture, 3D hydrogels were washed three times with 1× PBS, and 10 µL of 1× PBS was added over each hydrogel. A non-barcoded ITO glass slide was placed on top, and square-wave pulses (50 Hz, 200 µs pulse width, 1.0% duty cycle) were applied at 0–0.6 kV/cm for 1 min or at 0.4 kV/cm for 0–4 min. After 5 min, the buffer containing extracted RNA was collected, purified, and buffer-exchanged into nuclease-free water using an RNA Clean & Concentrator-5 kit (Zymo Research, R1013), and eluted in 10 µL.

RNA concentration was measured with the Qubit RNA HS Assay Kit (Thermo Fisher Scientific, Q32855), and the mass of extracted RNA was calculated as the measured concentration multiplied by the 10 µL elution volume. RNA extraction efficiency (%) was calculated as the extracted RNA mass divided by the estimated total RNA in each well, which was obtained by multiplying the number of cells per well by an assumed RNA content of 10 pg per cell.^42^ Because per-cell RNA content was assumed rather than measured, efficiencies are comparable across conditions but are not absolute. Non-electroporated samples (0 kV/cm) served as negative controls. Each condition was measured in three technical replicates, and viability was assessed 24 h after sampling as described below.

For repeated sampling, hydrogels were electroporated at 0.4 kV/cm for 1 min on days 1, 3, and 5 (D1, D3, and D5), and viability was imaged at baseline (D0) and immediately after each sampling event. Non-electroporated hydrogels were cultured in parallel as controls. Nanostraw-based sampling was performed as described previously^25^ at 0.4 kV/cm (20 Hz, 200 µs pulse width) for 1 min, with a 10 µm gap filled with 1× PBS between the nanostraws and ITO glass. Nanostraw extraction efficiency was calculated as described above.

### Extraction distribution assessment

To compare transcript representation in extracted RNA with that of whole-cell lysates, RNA extracted by electroporation (0.4 kV/cm, 1 min) was pooled from three hydrogels and purified as described above. Total RNA from separate unsampled hydrogels was extracted with TRIzol reagent (Thermo Fisher Scientific, 15596018), using GlycoBlue coprecipitant (Thermo Fisher Scientific, AM9515) to visualize the pellet. cDNA was synthesized with the iScript cDNA Synthesis Kit (Bio-Rad, 1708890) on a SimpliAmp thermal cycler (Thermo Fisher Scientific) according to the manufacturer’s instructions.

Each 10 µL qPCR reaction contained PowerUp SYBR Green Master Mix (Applied Biosystems, A25742), 1 ng of cDNA, and 500 nM each of forward and reverse primers (sequences in **Supplementary Table 1**). Reactions were run for 40 cycles on a QuantStudio 3 Real-Time PCR System (Applied Biosystems) using the manufacturer’s standard cycling conditions. ΔCt values were calculated relative to *18S rRNA* for *FAP*, *MITF*, *BRAF*, *ACTA2*, *HPRT1*, *MYC*, *PMEL*, *ACTB*, *HSPB1*, *YWHAZ*, *GAPDH*, *VIM*, *COL1A1*, *RPL13A*, and *EEF1A1*.

### Post-sampling gene expression assessment

Hydrogels were electroporated once (0.4 kV/cm, 1 min) and cultured for 48 h alongside non-electroporated controls. RNA was then extracted with TRIzol reagent and GlycoBlue coprecipitant as described above. cDNA synthesis and qPCR were performed as described above.

### Cell and tissue viability

Cell viability was assessed using a live/dead cell double staining kit (Sigma-Aldrich, 04511). Cells and hydrogels were incubated with calcein green AM (1 µM) and propidium iodide (1.5 µM) in DPBS containing calcium and magnesium for 15 min at 37 °C, and tissue slices for 1 h at 37 °C. Samples were imaged on an IX83 inverted microscope (Olympus). Live (calcein-positive) and dead (propidium iodide-positive) cells were counted in CellProfiler, FIJI, or manually for images with few cells. Viability was calculated for each replicate as live cells divided by total cells × 100, and averaged across replicates.

### Cell proliferation assay

Metabolic activity of tissue slices was measured with the CellTiter 96 AQueous One Solution Cell Proliferation Assay (MTS; Promega, G3580). Slices were incubated with 20 µL of reagent in 100 µL of medium for 1.5 h at 37 °C. Plates were then shaken on a GloMax Discover Microplate Reader (Promega, GM3000; linear motion, 10 s, 300 cycles per min, 2 mm diameter), and absorbance was measured at 490 nm. For the longitudinal study, absorbance of drug-treated slices was normalized to that of nontreated slices at each time point.

### Bulk RNA sequencing

Tissue slices from Patient 1 were collected 48 h after Live-ST sampling, alongside unsampled control slices from the same patient. RNA was extracted with TRIzol reagent and GlycoBlue coprecipitant as described above. cDNA was synthesized in a 20 µL reaction containing 1× Maxima RT buffer, 1 M betaine, 500 µM dNTPs, 1 U/µL RNase inhibitor, 5 µM biotin-oligo(dT)_30_, 1.25 µM template-switching oligonucleotide (sequences in **Supplementary Table 1**), and 50 U of Maxima H Minus reverse transcriptase (Thermo Fisher Scientific, EP0751) on a SimpliAmp thermal cycler for 30 min at 50 °C and 5 min at 85 °C. Because the oligo(dT) primer and the template-switching oligonucleotide carry the same 5′ handle, the cDNA had a template-switch primer site at both ends and was amplified by 12 cycles of PCR with the template-switch primer alone in 1× KAPA HiFi HotStart ReadyMix (Roche, KK2602). Amplified cDNA was purified sequentially with 0.8× and 0.6× solid-phase reversible immobilization (SPRI) beads. Libraries were prepared with the Nextera XT DNA Library Preparation Kit (Illumina, FC-131-1024), purified with 0.7× SPRI beads, and sequenced as described below. Read 2 of each bulk library was aligned as single-end reads to GRCh38 (primary assembly) with the GENCODE v45 annotation using STAR (v2.7.11b), which also counted uniquely mapped reads per gene (--quantMode GeneCounts). Strand-specific counts were used if they exceeded those of the opposite strand at least threefold; otherwise, unstranded counts were used. Gene counts were converted to counts per million (CPM) over all genes, and Pearson’s r was calculated on log10 CPM over genes detected in both samples (CPM > 0 in both), excluding ribosomal protein and mitochondrial genes.

### Flow cytometry

Melanoma tissue was washed three to four times in ice-cold PBS containing penicillin-streptomycin (1:50) to remove fat and debris, cut into 0.5–1 mm pieces with blades, and transferred to a 50 mL conical tube in pre-warmed digestion medium (10 mL per 0.2 g of tissue). Digestion medium consisted of DMEM without FBS supplemented with collagenase type 3 (1.5 mg/mL; Worthington Biochemical, LS004180), DNase I (0.1 mg/mL; Sigma-Aldrich, 4716728001), and Y-27632 ROCK inhibitor (1:1000; STEMCELL Technologies, 72307). Tissue was digested for 30 min at 37 °C on an orbital shaker with gentle vortexing halfway through. Digestion was stopped by adding 30–40 mL of complete medium (containing 10% FBS), and the suspension was filtered through a 70 µm cell strainer pre-wetted with 5 mL of complete medium. Cells were pelleted at 620 × *g* for 15 min, resuspended in complete medium, counted, and assessed for viability by acridine orange/propidium iodide staining. Cells were then washed twice in 10 mL of DPBS without calcium and magnesium (400 × *g*, 5 min; used for all washes), fixed in 5 mL of 4% paraformaldehyde in DPBS for 10 min at room temperature, and washed three times in DPBS. Fixed cells were frozen in 90% DMEM (containing 10% FBS) and 10% DMSO in a Mr. Frosty freezing container (Thermo Fisher Scientific) and stored at −80 °C. Cells frozen without fixation were fixed as above after thawing. Before staining, cells were washed three times in DPBS. Cells for intracellular staining were then permeabilized in 5 mL of 0.2% Tween 20 in 1% BSA in DPBS for 20 min at room temperature, and all cells were washed three times in 1% BSA in DPBS.

Cells were stained with Hoechst 33342 (Thermo Fisher Scientific, H3570) and with antibodies against SOX10, HMB45, CD45, CD3, CD19, CD11b, α-SMA, and CD31 (fluorophores, suppliers, and catalog numbers in Supplementary Table 1), with each marker stained in a separate sample alongside unstained and Hoechst-only controls. SOX10, CD45, α-SMA, and CD31 were detected with directly conjugated antibodies; HMB45, CD3, CD19, and CD11b were detected with unconjugated primary antibodies followed by Alexa Fluor 488- or Alexa Fluor Plus 488-conjugated secondary antibodies. Intracellular markers (SOX10, HMB45, and α-SMA) were stained in cell staining buffer (BioLegend, 420201) containing 0.1% saponin, with primary antibody incubation for 1.5 h at room temperature and gentle vortexing every 15 min, followed, where required, by secondary antibody incubation in the same buffer for 30 min at room temperature with gentle vortexing every 10 min. Surface markers were stained in the same way in cell staining buffer without saponin, with 30 min incubations for both primary and secondary antibodies. After each incubation, cells were washed three times with 1 mL of the same staining buffer, and stained cells were resuspended in 1 mL of cell staining buffer for acquisition.

Approximately 10,000 events per sample were acquired on a BD FACSymphony A3 Lite flow cytometer (BD Biosciences) at the Penn Cytomics and Cell Sorting Shared Resource Laboratory, University of Pennsylvania, and data were analyzed in FlowJo (v10.10.1; BD Biosciences). Cells were gated on forward scatter area (FSC-A) versus side scatter area (SSC-A) to exclude debris, followed by Hoechst-positive nucleated cells, and marker-positive gates were set from the unstained and Hoechst-only controls. Cell type proportions were defined as the percentages of tumor (SOX10-positive for the spatial mapping comparison; HMB45-positive for the drug-response study), α-SMA-positive (fibroblast), CD31-positive (endothelial), CD3-positive (T cell), CD19-positive (B cell), and CD11b-positive (myeloid) events, scaled to sum to 100% across the six cell types for comparison with Live-ST.

### ITO glass functionalization for spatial barcoding

ITO-coated glass was cut to 8.5 × 2.5 cm and sonicated sequentially in acetone, isopropanol, and DI water for 20 min each. Slides were rinsed with DI water and stored in DI water, protected from light, for at least 3 days before silanization. Slides were dried with compressed air and treated with oxygen plasma (Harrick Plasma, 4768581; 0.4–0.7 Torr, high setting) for 30 s, followed by 30 s under vacuum. Plasma-treated slides were immediately immersed in 3% (v/v) 3-aminopropyltriethoxysilane (APTES; Sigma-Aldrich, 440140) in acetone and incubated for 2 h at room temperature under nitrogen. Slides were washed twice with acetone, dried with compressed air, and stored dry at 4 °C overnight.

APTES-modified slides were then covered with 2.5% (v/v) glutaraldehyde (Sigma-Aldrich, G5882) in 1× PBS and incubated for 2 h at room temperature. Slides were rinsed three times with 1× PBS and once with molecular-grade water, dried with compressed air, and stored dry at 4 °C, protected from light, until barcoding.

### Microfluidic spatial barcoding on ITO glass

Spatial barcodes were generated by combinatorial deterministic barcoding with PDMS microfluidic devices, following previously described approaches.^4,32,33^ Each device contained 50 parallel channels (50 µm wide and 50 µm high, with 50 µm spacing) and was fabricated by soft lithography. All oligonucleotides were from IDT (sequences in Supplementary Table 1).

#### Barcode A immobilization

Amine-modified barcode A oligonucleotides (100 µM in RNase-free water) were mixed with 0.167 M MES buffer (pH 6.5) at a 2:3 ratio (final oligonucleotide concentration 40 µM). The first device, with channels along the x-axis, was placed on the APTES-glutaraldehyde-modified slide, and 5 µL of each of the 50 barcode A solutions was loaded into its channel and drawn through by vacuum (−2 psi until the channels filled, then −0.8 psi for 1 min). Slides were incubated for 4 h at room temperature in a humidified chamber, protected from light. The device was removed, and slides were washed once with 2× SSC containing 0.1% SDS for 10 min, once with 0.2× SSC for 2 min, and once with molecular-grade water, and dried with compressed air.

#### Barcode B ligation

Each 5′-phosphorylated barcode B oligonucleotide, containing a unique molecular identifier (UMI; 12 nt) and a poly(T)_28_ capture sequence, was annealed to a linker oligonucleotide at 50 µM each in annealing buffer (10 mM Tris-HCl pH 8.0, 20 mM NaCl) by heating to 95 °C for 1 min and cooling to 20 °C at 0.1 °C/s. Annealed barcode B–linker duplexes were mixed 1:1 with ligation mix to final concentrations of 25 µM duplex, 1× T4 DNA ligase buffer, 40 U/µL T4 DNA ligase (NEB, M0202S), and 200 µg/mL BSA. The second device, with channels along the y-axis, was placed orthogonally on the slide, and 5 µL of each of the 50 ligation mixtures was loaded and drawn through by vacuum as above. Ligation proceeded for 2 h at room temperature in a humidified chamber, protected from light. Slides were washed as above, dried, and stored dry at 4 °C, protected from light. This produced a 50 × 50 grid of 2,500 uniquely barcoded spots within a 5 × 5 mm area.

#### Barcode validation

To validate each reaction, barcode A (40 µM) or the barcode B ligation mixture was applied in bulk (40 µL under a HybriSlip) to functionalized slides and processed as above. Fluorescently labeled barcode B (FAM) was detected only on barcode A-functionalized slides and not on control slides, and crossed channels of labeled barcode A (Cy5) and barcode B (FAM) produced the expected checkerboard pattern. Slides were imaged on the IX83 microscope.

### RNA spatial preservation in fixed A549 cells with optical readout

Capture slides for optical readout were functionalized with amine-modified poly(T) capture probes (sequence in Supplementary Table 1; adapted from Enhanced ELectric Fluorescence in situ Hybridization) using the APTES/glutaraldehyde chemistry described above.^24^ For cell patterning, ITO glass slides were cleaned with ethanol and DI water, treated with oxygen plasma, and coated with undiluted poly-L-lysine solution (0.01%; Sigma-Aldrich, P4707) for 1 h at room temperature. Acrylic “T”-shaped wells were laser-cut (1/8 inch thick; Rapid Prototyping Laboratory), placed within a PDMS reservoir on ITO glass, and seeded with A549 cells. After 24 h, cells were fixed in 70% ethanol in molecular-grade water overnight at 4 °C (a non-cross-linking fixative, so that RNA could be released by electroporation), washed three times in DPBS, and the well was removed; nuclei were stained with Hoechst 33342 and imaged before and after stamping.

The capture slide was placed over the fixed cells using the stamping and electroporation setup described above, and pulses were applied at 2 kV/cm for 30 s (50 Hz, 200 µs pulse width, 1.0% duty cycle); non-electroporated samples served as controls. Captured RNA was reverse transcribed on the slide with Maxima H Minus reverse transcriptase as described above; single-stranded DNA (unextended capture probes) was digested with exonuclease I (3′→5′; NEB, M0293L) for 45 min at 37 °C, and the remaining RNA-cDNA hybrids were stained with SYBR Gold (Thermo Fisher Scientific, S11494; 1:1000) in 2× SSC and imaged on the IX83 microscope. To validate the optical workflow, 0 or 1,000 ng of purified A549 total RNA was incubated on capture slides for 30 min at room temperature in 1× PBS and processed identically.

#### Image analysis

SYBR Gold (cDNA) and Hoechst (cells) images (4×; 620 pixels per mm from the scale bar) were analyzed in Python (OpenCV, SciPy). The Hoechst image was registered to the SYBR Gold image by translation, maximizing the overlap of the two masks. Each channel was smoothed with a Gaussian filter (σ = 4 µm) and thresholded by Otsu’s method, and the masks were closed morphologically (40 µm); only the largest connected component was kept, within a field excluding the channel walls. Spatial agreement was quantified as the Dice coefficient between the two masks and as the fraction of the cDNA mask area lying within the cell mask. Lateral spread was measured from an edge profile: each pixel was assigned its signed distance from the cell-mask boundary, and the mean SYBR Gold intensity was calculated in 2-µm distance bins, excluding pixels within 300 µm of the field border. The profile was scaled from the background level (150–250 µm outside the boundary) to the level inside the cells (40–100 µm inside), and lateral spread was taken as the distance at which the cDNA signal fell to half. Because Hoechst stains only nuclei, this distance includes cytoplasm beyond the outermost nuclei and is an upper bound. As a sensitivity check, lateral spread was recalculated for all nine combinations of three inside (20–60, 40–100, and 80–150 µm inside the boundary) and three background (100– 200, 150–250, and 200–250 µm outside) reference windows, and the largest value is reported as an upper limit alongside the main estimate.

### Live Spatial Transcriptomics pipeline

#### Sampling

Live cells or tissue slices were placed on the tissue-supporting ITO slide (cathode) in 10–20 µL of 1× PBS, and the barcoded capture slide (anode) was lowered with the dovetail stage until contact was detected by the load cell. Square-wave pulses (0.4 kV/cm, 50 Hz, 200 µs pulse width, 1.0% duty cycle) were applied for 1 min for live cells or 10 s for live tissue slices; the shorter duration was used for tissue because longer contact caused the tissue to adhere to the barcoded capture slide and strip the barcodes from the glass. After 30 s, the capture slide was removed, and the cells or tissue were returned to culture.

#### RT and second-strand synthesis

After sampling, the capture area was kept hydrated throughout and washed once with 0.1× SSC in molecular-grade water. Captured RNA was reverse transcribed on the slide in 40 µL containing 1× Maxima RT buffer, 2 U/µL RNase inhibitor, 0.2 mg/mL BSA, 1 mM dNTPs, 2.5 µM template-switching oligonucleotide, and 10 U/µL Maxima H Minus reverse transcriptase at 42 °C for 90 min in a humidified chamber. The slide was washed with 0.1× SSC, incubated with 0.08 M KOH for 5 min to remove RNA, and washed again with 0.1× SSC. Second-strand synthesis was performed on the slide in 40 µL containing 1× isothermal amplification buffer, 2.5 µM template-switch primer, 4 mM MgSO_4_, 0.32 U/µL *Bst* 2.0 DNA polymerase (NEB, M0537S), and 1 mM dNTPs at 65 °C for 15 min, followed by a 0.1× SSC wash.

#### cDNA release and amplification

Second-strand products were released by incubating with 35 µL of 0.08 M KOH for 10 min at room temperature with pipetting every 5 min, collected, and neutralized with 5 µL of 1 M Tris-HCl (pH 7.0). Single-stranded DNA was purified with a DNA Clean & Concentrator kit-25 (Zymo Research, R1018; 7:1 binding buffer ratio) and eluted in 24 µL of nuclease-free water. cDNA was amplified in a 50 µL reaction containing 1× KAPA HiFi HotStart ReadyMix and 1 µM each of the template-switch primer and primer 1, with cycling at 95 °C for 3 min; 16 cycles of 98 °C for 20 s, 65 °C for 30 s, and 72 °C for 2 min; and 72 °C for 5 min. Amplified cDNA was purified with 0.8× SPRI beads, eluted in 20 µL of nuclease-free water, and assessed on a TapeStation D5000 assay (Agilent).

#### Library preparation and sequencing

Sequencing libraries were prepared from 1 ng of cDNA (quantified by Qubit) with the Nextera XT DNA Library Preparation Kit using 0.5 µM each of the P5-N501 primer and an indexed P7 primer (N7xx; sequences in Supplementary Table 1), purified with 0.7× SPRI beads, eluted in 15 µL of nuclease-free water, and assessed on a TapeStation D5000 assay. Libraries were sequenced as described below.

### RNA spatial preservation in live A549 cells with sequencing readout

Four PDMS reservoirs (1 mm diameter) were placed within the 5 × 5 mm barcoded region on poly-L-lysine-coated ITO glass (prepared as described above) and seeded with A549 cells. After 24 h, the reservoirs were removed, and the patterned cells were imaged by bright-field microscopy and processed by Live-ST as described above. Viability was assessed 24 h after sampling in the four patterned regions as described above, and sequencing data were analyzed as described below.

### Spatial mapping of living tissue

A 5 mm melanoma slice from Patient 2 was marked with a 1 mm biopsy punch at one edge as a positional landmark and processed by Live-ST as described above. Data were analyzed as described below.

### Longitudinal profiling of drug responses in the tumor-immune microenvironment

*BRAF* V600E-mutant melanoma tissue from Patient 4 was cored with a 6 mm biopsy punch, sectioned into 400 µm slices, and marked with a 1 mm biopsy punch at the tissue edge for alignment. Slices were profiled by Live-ST at D-1, D1, and D3. Trametinib (VWR, 77609-120) and dabrafenib (VWR, MSPP-HY14660) were added at D0 at 10 nM (trametinib) and 100 nM (dabrafenib) to two slices (Treated 1 and 2), and two slices were maintained without drug (Nontreated 1 and 2). Concurrently, additional drug-treated and nontreated slices from the same patient were cultured with or without Live-ST sampling as viability controls. A slice was submitted to the Department of Pathology and Laboratory Medicine for H&E staining. Remaining tissue from the same patient was dissociated and analyzed by flow cytometry as described above.

### Spatial sequencing and read processing

Spatial and bulk libraries were sequenced on an AVITI system (Element Biosciences) with paired-end reads (Read 1, 99 nt; Read 2, 199 nt) using the AVITI 2 × 150 Sequencing Kit Cloudbreak FS. Reads were demultiplexed by sample index with bases2fastq (v2.3.0). In spatial libraries, Read 1 contains barcode A, barcode B, and the UMI; Read 2 contains the cDNA.

Spatial barcodes and UMIs were extracted from Read 1 with custom Python scripts. Read 1 consists of an 8-nt barcode A, an 18-nt linker, an 8-nt barcode B, and a 12-nt UMI. Barcodes A and B were each matched to the 50 barcodes of their set, allowing one mismatch, which assigned the read pair to one spot of the 50 × 50 array (2,500 spots). Reads whose barcodes started one base late were also assigned, and read pairs without both a matching barcode A and barcode B were discarded.

Read 2 was trimmed in two steps. First, the template-switching oligonucleotide at the start of the read (at least its last 12 nt, with the Gs that follow it) was removed. Second, the read was trimmed at the first poly(A) stretch (8 A), TruSeq or Nextera adapter sequence, or read-through into the capture oligonucleotide, identified by the reverse complement of the Read 1 linker (allowing one mismatch, including a partial linker at the end of the read) or by the reverse-complemented barcode B and barcode A sequences in their expected arrangement. Reads shorter than 10% of their original length (20 nt) after trimming were discarded.

Distinct Read 2 sequences were aligned to GRCh38 (primary assembly) with the GENCODE v45 annotation using STAR with permissive settings (--outFilterScoreMinOverLread 0.1, -- outFilterMatchNminOverLread 0.45, --outFilterMismatchNmax 3, --alignSJDBoverhangMin 1). Only uniquely mapped reads were kept (--outFilterMultimapNmax 1). A read was assigned to a gene if it overlapped an annotated exon of that gene on the same strand. Uniquely mapped reads outside exons (intronic, intergenic, or antisense) were kept as molecules without a gene, so gene counts include exonic reads only.

Within each spot, reads were grouped by UMI. A UMI that differed by one base from a UMI of the same gene with at least 2*n* − 1 reads, where *n* is its own read count, was merged into it (the directional method of UMI-tools), and each molecule was assigned the gene supported by most of its reads. Two counts were kept per spot: all uniquely mapped molecules, used for tissue detection and the UMI maps, and gene-assigned molecules, used for all other analyses. Read pairs per sample and the fractions of distinct Read 2 sequences that mapped uniquely, mapped to multiple loci, or did not map were taken from the read-processing log and the STAR summary statistics.

### Spatial transcriptome analysis

#### Cell-pattern test array

The live A549 cell array was processed with the same read processing, alignment, and UMI counting as above. The bright-field image (grayscale; cells dark) was mirrored and rotated to the orientation of the array, scaled from its 200 µm scale bar (610 pixels per mm; 10 spots per mm), centered on the 50 × 50 grid, and sampled at each spot center by bilinear interpolation. The sampled image was inverted so that cells had high values, background (inverted intensity < 60 of 255) was set to 0, and the result was smoothed with a Gaussian filter (σ = 1.1 spots) and square-root scaled to 0–2 for display. The UMI map (all uniquely mapped molecules per spot) was transformed to log(UMI + 1), smoothed with a Gaussian filter (σ = 2.5 spots), and z-scored over the array. The image and UMI maps were compared by their contours at 1.0 (image scale and UMI z-score) and by line profiles at barcode coordinates X = 15 and 35 and Y = 15 and 35. UMIs and genes per spot were counted from gene-assigned molecules over all spots with at least one such molecule.

#### Tissue detection and cell type mapping

Spots with at least one uniquely mapped molecule passed quality control (at least 25 in the spatial mapping experiment), and log(UMI + 1) was z-scored over the passing spots of each sample, for the background threshold below. Cell types were mapped with cell2location^43^ (v0.1.5; scvi-tools v1.4.3; random seed 0) using a melanoma single-cell RNA sequencing (scRNA-seq) reference (GSE115978^34^; 7,186 cells; raw counts) with the TISCH2 major-lineage labels^44^, grouped into tumor cells (malignant), T cells (conventional CD4+, exhausted CD8+, and proliferating T cells, and natural killer cells), B cells, monocytes/macrophages, fibroblasts, and endothelial cells. Reference signatures were estimated with the cell2location regression model (150 epochs) and cell type abundances with the spatial model (1,000 epochs; N_cells_per_location = 3; detection_alpha = 20), over the genes shared by the reference and the spatial data, from gene-assigned UMI counts. Spots that passed quality control and had at least one UMI in these genes were mapped, and each was assigned its most abundant cell type (posterior mean of 1,000 posterior samples). Per-spot pie charts show each mapped spot’s four most abundant cell types, rescaled to sum to 100%. Spots with a z-scored log(UMI + 1) below a background threshold (−4 in the drug-response study; −1.7 in the spatial mapping experiment) were left unclassified. Spots assigned a cell type are referred to as tissue spots, and cell type proportions are fractions of tissue spots, not of cells.

#### Tissue image registration

In the spatial mapping experiment, the bright-field image of the tissue slice was registered to the spot grid: it was scaled from its 1 mm scale bar (612 pixels per mm, measured in FIJI; 10 spots per mm), mirrored and rotated to the orientation of the array, and translated by eye until the tissue matched the UMI map. The UMI map in **Fig. 3j** instead shows the z-score of log(UMI + 1) over all 2,500 spots, with spots that failed quality control in grey.

#### Drug-response study

Each tissue was sampled on the same array at all three time points, so array coordinates refer to the same location in a tissue over time. Tissues were the biological replicates (*n* = 2 per group), and spots were not treated as replicates. Each sample’s pseudobulk profile was the sum of gene-assigned UMI counts over its tissue spots.

#### Quality control and reproducibility

UMIs and detected genes per spot were counted from gene-assigned molecules over all spots with data. Pseudobulk expression was calculated as CPM, and Pearson’s *r* was calculated on log10 CPM over genes detected in both samples (CPM > 0 in both), between the time points of each tissue. Ribosomal protein genes (exact RPL/RPS gene symbols, so that RPS6K kinases were kept) and mitochondrial genes were excluded from the pseudobulk correlations.

#### Pathway activity over time

The ERK output signature comprised the 181 genes downregulated by at least three of four MEK inhibitors (trametinib, cobimetinib, selumetinib, and PD-0325901) in the LINCS L1000 consensus signatures (Enrichr library LINCS_L1000_Chem_Pert_Consensus_Sigs). Glycolysis and TNF-α signaling via NF-κB gene sets were taken from MSigDB Hallmark 2020; the TNF-α signaling set excluded genes shared with the ERK output signature. Genes with at least 5 UMIs in at least 2 of the 12 pseudobulks were considered detected, and only detected genes were scored (ERK output, 93 genes; glycolysis, 111; TNF-α signaling, 82). For each tissue, the three pseudobulks were randomly downsampled together (binomial thinning, single draw, fixed seed) to the smallest of their totals, so that the time points of a tissue were equally deep; tissues were not matched to each other. Each gene’s change from D-1 was log_2_((UMIs at D1 or D3 + 1)/(UMIs at D-1 + 1)). A pathway score was the mean change of its genes minus the mean change of all detected genes, so that a random gene set scores about 0; scores are 0 at D-1 by definition. Group values are the mean of the two tissues. As a reference for chance, the central 95% range of the scores of 1,000 random sets of detected genes of the same size was calculated.

#### ERK output gene changes

For each ERK output gene with at least 5 UMIs at D-1 and D3 combined, after downsampling, in every tissue, the log_2_ change from D-1 to D3 was calculated for each tissue as above.

#### Tumor-spot comparison

To assess the effect of cell type composition, pathway scores were also calculated from pseudobulks of tumor spots only (spots whose most abundant cell type was tumor cells). For the comparison with all tissue spots, both pseudobulks of a tissue were downsampled to the depth of its smallest tumor-spot pseudobulk; values are the mean ± SD of 20 downsampling draws.

#### Spatial maps of pathway activity

Because each spot captures too few molecules to score a gene set, pathway activity was mapped over 7 × 7-spot windows of tissue spots centered on each tissue spot. In each window, the observed count O was the window’s UMIs in the pathway’s genes, and the expected count E was the window’s total UMIs multiplied by the pathway’s share of UMIs (the mean over the 12 tissue-spot pseudobulks, each sample weighted equally). The score log_2_((O + 0.5)/(E + 0.5)) was corrected by subtracting the mean score of 50 control gene sets in the same window, each replacing every pathway gene with a random detected gene of similar abundance (the same one of 25 bins of study-wide UMI share), as in Seurat’s AddModuleScore. A score of 0 means that the pathway is at its study-wide average share. Windows with E < 1 are not shown. Pathway genes were the detected genes defined above. Color scales are symmetric about 0 and span the 90th percentile of absolute scores across the maps shown.

#### Most abundant genes

For each time point, the four pseudobulks were downsampled to the smallest of their totals (binomial thinning, fixed seed), and expression was calculated as log(CPM + 1); a group’s value averaged the CPM of its two tissues. The 15 most abundant genes per group were selected and grouped by whether they were among the most abundant in treated tissues only, nontreated tissues only, or both. The list was topped up to 30 genes with the next most abundant genes detected in both groups, ranked by mean log(CPM + 1). Genes tied at the cutoff were all kept. Unannotated genes (Ensembl IDs, LOC and LINC), ribosomal protein genes, and mitochondrial genes were excluded.

#### GO enrichment

GO enrichment was performed with GSEApy (v1.3.1; enrichr function) against the GO Biological Process 2021 library, with all genes in the library as background. For each group and time point, the 50 most abundant genes (genes tied at the cutoff left out) were tested with a one-sided hypergeometric test with Benjamini–Hochberg correction; terms with adjusted *P* < 0.05 were considered significant. For each group, the 10 terms with the lowest adjusted *P* were shown, topped up to 20 terms per time point.

### Statistics

Unless stated otherwise, data are shown as mean ± SD, and *n* is defined in each figure legend. Viability in the repeated-sampling experiment was compared between electroporated samples and non-electroporated controls with an unpaired two-tailed t-test on the mean viability at each time point. ΔCt values in the extraction distribution and post-sampling gene expression assessments were compared with unpaired two-tailed t-tests across targets. These tests were performed in GraphPad Prism (v11.1.0, GraphPad Software), and significance was set at *P* < 0.05. In the drug-response study, *n* is the number of tissues (biological replicates; two per group). With two tissues per group, pathway scores were not tested statistically: each tissue is shown with the group mean, and random gene sets of the same size give a reference for chance. Pearson correlations and comparisons of cell type composition between Live-ST and flow cytometry are descriptive and were not tested statistically. GO enrichment *P* values were adjusted for multiple testing with the Benjamini–Hochberg method. Random downsampling used a fixed seed. Unless stated otherwise, log denotes the natural logarithm. All other analyses were performed in Python (v3.13).

## Acknowledgements

The data for this manuscript were generated in the Penn Cytomics and Cell Sorting Shared Resource Laboratory at the University of Pennsylvania (RRID:SCR_022376). Penn Cytomics is partially supported by the Abramson Cancer Center NCI Grant (P30 CA016520). We thank the University of Pennsylvania Libraries’ Holman Biotech Commons for fabrication of the custom slide holders and tissue stamper, and the Rapid Prototyping Laboratory for the laser-cut acrylic wells. We also thank the Department of Pathology and Laboratory Medicine at the Hospital of the University of Pennsylvania for providing patient tissue samples and performing H&E staining.

## Author information

These authors contributed equally: David E. Reynolds and Sangmo Lee

---

**Extended Data Figure 1:**
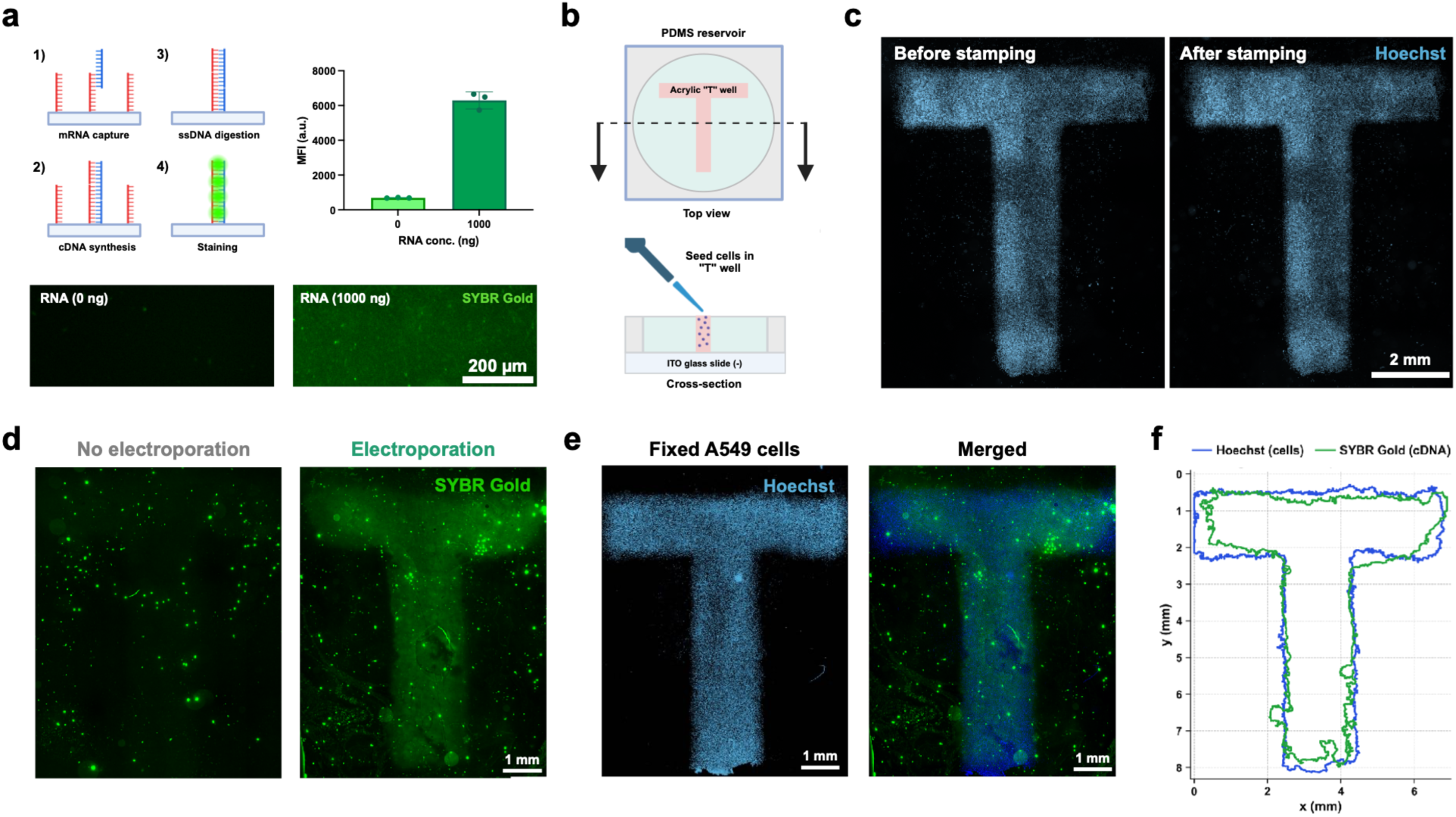
Spatial preservation of captured RNA. (**a**) Optical capture workflow (left): (1) mRNA capture by amine-modified poly(T) capture probes on ITO glass, (2) cDNA synthesis by reverse transcription, (3) exonuclease (3′→5′) digestion of single-stranded DNA (unextended capture probes), and (4) SYBR Gold staining of RNA-cDNA hybrids. Mean fluorescence intensity (MFI) after incubation with 0 or 1,000 ng of purified A549 RNA (right), with representative images (bottom). (**b**) Patterning of A549 cells in an acrylic “T”-shaped well within a PDMS reservoir on ITO glass (top and cross-section views). Created with BioRender.com. (**c**) Hoechst-stained patterned cells before and after stamping. (**d**) SYBR Gold-stained cDNA on the capture slide without and with electroporation. (**e**) Hoechst-stained fixed cells and the merged image with captured cDNA. (**f**) Outlines of the cell (Hoechst, blue) and cDNA (SYBR Gold, green) masks, generated by Gaussian smoothing, Otsu thresholding, and morphological closing. Dice coefficient = 0.88; 96% of the cDNA signal area lies within the cell-covered area, and cDNA signal falls to half its intensity within ∼19 µm of the cell boundary. Data in (**a**) are shown as mean ± SD (*n* = 3 fields of view per condition). Scale bars: 200 µm (**a**), 2 mm (**c**), 1 mm (**d**,**e**).

**Extended Data Figure 2:**
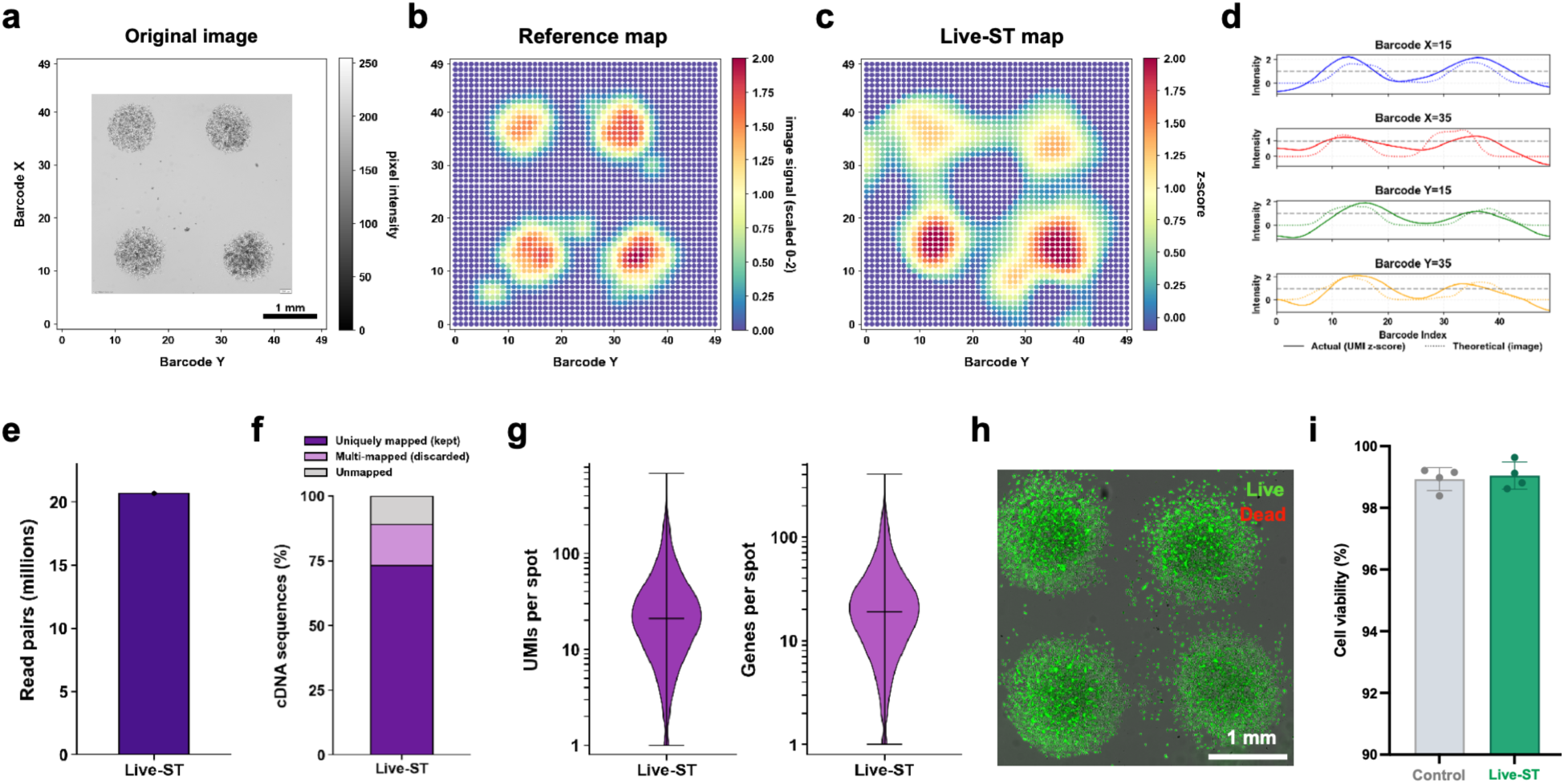
Spatial preservation of captured RNA with living cells. (**a**) Bright-field image of live A549 cells patterned in four 1 mm regions within the 5 × 5 mm barcoded area, shown on barcode coordinates. (**b**) Reference map derived from the image in (a), scaled from 0 to 2. (**c**) Live-ST map of UMI counts per spot (log(UMI + 1), smoothed and z-scored). (**d**) Line profiles of theoretical (image-derived; dotted) and measured (UMI z-score; solid) intensity along barcode coordinates X = 15 and 35 and Y = 15 and 35. (**e**) Number of read pairs. (**f**) Fraction of cDNA sequences that were uniquely mapped, multi-mapped (discarded), or unmapped. (**g**) UMIs (left) and genes (right) per spot (median ∼20 each). (**h**) Live/Dead staining 24 h after Live-ST sampling. Live cells are shown in green and dead cells in red. (**i**) Cell viability of control and Live-ST-sampled cells 24 h after sampling. Data in (**i**) are shown as mean ± SD (*n* = 4 patterned cell regions per condition). Scale bars: 1 mm (**a**,**h**).

**Extended Data Figure 3:**
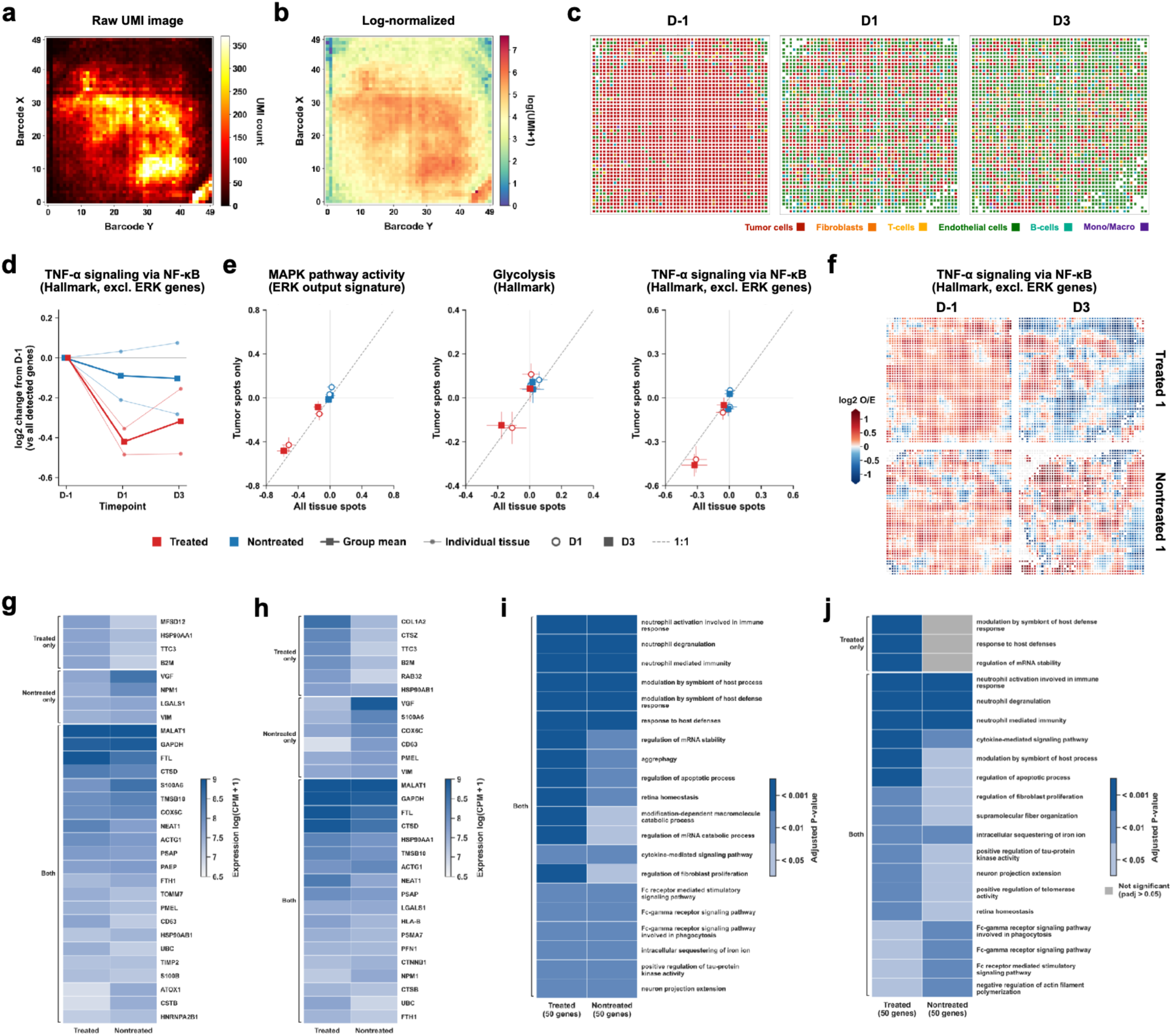
Longitudinal profiling of drug responses, additional analysis. (**a**,**b**) Raw UMI counts (**a**) and log-normalized expression, log(UMI + 1) (**b**), per spot in Treated 1 one day before treatment (D-1). (**c**) Spatial maps of the dominant cell type in each spot in Treated 1 at D-1, D1, and D3. (**d**) Change from D-1 in the TNF-α signaling via NF-κB module score (Hallmark gene set excluding ERK output genes), shown as in **Figure 4g**. (**e**) Change from D-1 in ERK output (left), glycolysis (middle), and TNF-α signaling via NF-κB (right) module scores computed using tumor-dominant spots only versus all tissue spots, for treated (red) and nontreated (blue) tissues at D1 (open symbols) and D3 (filled symbols). The dashed line indicates 1:1. (**f**) Spatial maps of TNF-α signaling via NF-κB (log_2_ observed/expected) at D-1 and D3 in Treated 1 and Nontreated 1. (**g**,**h**) Expression (log(CPM + 1)) of the most abundant transcripts in treated and nontreated tissues at D1 (**g**) and D3 (**h**), grouped by whether they were among the most abundant in treated tissue only, nontreated tissue only, or both. (**i**,**j**) GO biological process terms enriched among the 50 most abundant genes at D1 (**i**) and D3 (**j**), colored by adjusted *P* value; grey indicates terms that were not significant (adjusted *P* > 0.05). In (**e**), each tissue’s all-spot and tumor-spot pseudobulks were downsampled to the same depth (that of its smallest tumor-spot pseudobulk), so all-spot values differ from those in (**d**) and **Figure 4g**; points show the mean of 20 random downsamplings and error bars the SD across them.

## References

1. Jain, S. & Eadon, M. T. Spatial transcriptomics in health and disease. Nat Rev Nephrol 20, 659–671 (2024).

2. Rao, A., Barkley, D., França, G. S. & Yanai, I. Exploring tissue architecture using spatial transcriptomics. Nature 596, 211–220 (2021).

3. Ståhl, P. L., Salmén, F., Vickovic, S., Lundmark, A., Navarro, J. F., Magnusson, J., Giacomello, S., Asp, M., Westholm, J. O., Huss, M., Mollbrink, A., Linnarsson, S., Codeluppi, S., Borg, Å., Pontén, F., Costea, P. I., Sahlén, P., Mulder, J., Bergmann, O., Lundeberg, J. & Frisén, J. Visualization and analysis of gene expression in tissue sections by spatial transcriptomics. Science 353, 78–82 (2016).

4. Liu, Y., Yang, M., Deng, Y., Su, G., Enninful, A., Guo, C. C., Tebaldi, T., Zhang, D., Kim, D., Bai, Z., Norris, E., Pan, A., Li, J., Xiao, Y., Halene, S. & Fan, R. High-Spatial-Resolution Multi-Omics Sequencing via Deterministic Barcoding in Tissue. Cell 183, 1665–1681.e18 (2020).

5. Rodriques, S. G., Stickels, R. R., Goeva, A., Martin, C. A., Murray, E., Vanderburg, C. R., Welch, J., Chen, L. M., Chen, F. & Macosko, E. Z. Slide-seq: A scalable technology for measuring genome-wide expression at high spatial resolution. Science 363, 1463–1467 (2019).

6. Tian, L., Chen, F. & Macosko, E. Z. The expanding vistas of spatial transcriptomics. Nat Biotechnol 41, 773–782 (2023).

7. Moses, L. & Pachter, L. Museum of spatial transcriptomics. Nat Methods 19, 534–546 (2022).

8. Reynolds, D. E., Roh, Y. H., Oh, D., Vallapureddy, P., Fan, R. & Ko, J. Temporal and spatial omics technologies for 4D profiling. Nat Methods 22, 1408–1419 (2025).

9. Chen, W., Guillaume-Gentil, O., Rainer, P. Y., Gäbelein, C. G., Saelens, W., Gardeux, V., Klaeger, A., Dainese, R., Zachara, M., Zambelli, T., Vorholt, J. A. & Deplancke, B. Live-seq enables temporal transcriptomic recording of single cells. Nature 608, 733–740 (2022).

10. Guillaume-Gentil, O., Grindberg, R. V., Kooger, R., Dorwling-Carter, L., Martinez, V., Ossola, D., Pilhofer, M., Zambelli, T. & Vorholt, J. A. Tunable Single-Cell Extraction for Molecular Analyses. Cell 166, 506–516 (2016).

11. Ji, X., Fang, P., Wan, Y., Zhao, X., Xiong, C., Guo, F., Yang, Q., Yan-Do, R., Qi, L., Xu, C., Wang, X., Huang, L., Zhang, W., Wang, Z., Ke, J. & Shi, P. Sequencing-free spatial profiling of post-transcriptional regulation in fresh tissues using nanoneedle arrays. Nat Biomed Eng (2026).

12. Wang, Z., Qi, L., Yang, Y., Lu, M., Xie, K., Zhao, X., Cheung, E. H. C., Wang, Y., Jiang, X., Zhang, W., Huang, L., Wang, X. & Shi, P. High-throughput intracellular biopsy of microRNAs for dissecting the temporal dynamics of cellular heterogeneity. Sci Adv 6, eaba4971 (2020).

13. Reynolds, D. E., Roh, Y. H., Chintapula, U., Huynh, E., Vallapureddy, P., Tran, H. H., Lee, D., Allen, M. G., Xu, X. & Ko, J. Vertically Aligned Nanowires for Longitudinal Intracellular Sampling. ACS Nano 19, 13073–13083 (2025).

14. Zhao, X., Ji, X., Qu, J., Xie, K., Wang, Z., Fang, P., Wang, Y., Wan, Y., Yang, Y., Zhang, W. & Shi, P. Sequencing-free Analysis of Multiple Methylations on Gene-Specific mRNAs. J Am Chem Soc 144, 6010–6018 (2022).

15. Xia, C., Colognori, D., Jiang, X. S., Xu, K. & Doudna, J. A. Single-molecule live-cell RNA imaging with CRISPR-Csm. Nat Biotechnol 43, 2023–2030 (2025).

16. Wang, H., Nakamura, M., Abbott, T. R., Zhao, D., Luo, K., Yu, C., Nguyen, C. M., Lo, A., Daley, T. P., La Russa, M., Liu, Y. & Qi, L. S. CRISPR-mediated live imaging of genome editing and transcription. Science 365, 1301–1305 (2019).

17. Legnini, I., Emmenegger, L., Zappulo, A., Rybak-Wolf, A., Wurmus, R., Martinez, A. O., Jara, C. C., Boltengagen, A., Hessler, T., Mastrobuoni, G., Kempa, S., Zinzen, R., Woehler, A. & Rajewsky, N. Spatiotemporal, optogenetic control of gene expression in organoids. Nat Methods 20, 1544–1552 (2023).

18. Qiu, Q., Zhang, H., Xia, Z., Gao, W., Leu, J. I., Liang, D., Li, Y., Li, F., Su, Y., Feierman, E., Van Horn, E., Ming, G.-L., Korb, E., Song, H., Zhou, Z. & Wu, H. Spatial mapping of RNA turnover kinetics in the mouse brain. Nat Neurosci (2026).

19. Ren, J., Zhou, H., Zeng, H., Wang, C. K., Huang, J., Qiu, X., Sui, X., Li, Q., Wu, X., Lin, Z., Lo, J. A., Maher, K., He, Y., Tang, X., Lam, J., Chen, H., Li, B., Fisher, D. E., Liu, J. & Wang, X. Spatiotemporally resolved transcriptomics reveals the subcellular RNA kinetic landscape. Nat Methods 20, 695–705 (2023).

20. Chen, A., Liao, S., Cheng, M., Ma, K., Wu, L., Lai, Y., Qiu, X., Yang, J., Xu, J., Hao, S., Wang, X., Lu, H., Chen, X., Liu, X., Huang, X., Li, Z., Hong, Y., Jiang, Y., Peng, J., Liu, S., Shen, M., Liu, C., Li, Q., Yuan, Y., Wei, X., Zheng, H., Feng, W., Wang, Z., Liu, Y., Wang, Z., Yang, Y., Xiang, H., Han, L., Qin, B., Guo, P., Lai, G., Muñoz-Cánoves, P., Maxwell, P. H., Thiery, J. P., Wu, Q.-F., Zhao, F., Chen, B., Li, M., Dai, X., Wang, S., Kuang, H., Hui, J., Wang, L., Fei, J.-F., Wang, O., Wei, X., Lu, H., Wang, B., Liu, S., Gu, Y., Ni, M., Zhang, W., Mu, F., Yin, Y., Yang, H., Lisby, M., Cornall, R. J., Mulder, J., Uhlén, M., Esteban, M. A., Li, Y., Liu, L., Xu, X. & Wang, J. Spatiotemporal transcriptomic atlas of mouse organogenesis using DNA nanoball-patterned arrays. Cell 185, 1777–1792.e21 (2022).

21. Sampath Kumar, A., Tian, L., Bolondi, A., Hernández, A. A., Stickels, R., Kretzmer, H., Murray, E., Wittler, L., Walther, M., Barakat, G., Haut, L., Elkabetz, Y., Macosko, E. Z., Guignard, L., Chen, F. & Meissner, A. Spatiotemporal transcriptomic maps of whole mouse embryos at the onset of organogenesis. Nat Genet 55, 1176–1185 (2023).

22. Chow, K.-H. K., Budde, M. W., Granados, A. A., Cabrera, M., Yoon, S., Cho, S., Huang, T.-H., Koulena, N., Frieda, K. L., Cai, L., Lois, C. & Elowitz, M. B. Imaging cell lineage with a synthetic digital recording system. Science 372, eabb3099 (2021).

23. Ratz, M., von Berlin, L., Larsson, L., Martin, M., Westholm, J. O., La Manno, G., Lundeberg, J. & Frisén, J. Clonal relations in the mouse brain revealed by single-cell and spatial transcriptomics. Nat Neurosci 25, 285–294 (2022).

24. Borm, L. E., Mossi Albiach, A., Mannens, C. C. A., Janusauskas, J., Özgün, C., Fernández-García, D., Hodge, R., Castillo, F., Hedin, C. R. H., Villablanca, E. J., Uhlén, P., Lein, E. S., Codeluppi, S. & Linnarsson, S. Scalable in situ single-cell profiling by electrophoretic capture of mRNA using EEL FISH. Nat Biotechnol 41, 222–231 (2023).

25. Cao, Y., Hjort, M., Chen, H., Birey, F., Leal-Ortiz, S. A., Han, C. M., Santiago, J. G., Paşca, S. P., Wu, J. C. & Melosh, N. A. Nondestructive nanostraw intracellular sampling for longitudinal cell monitoring. Proc Natl Acad Sci U S A 114, E1866–E1874 (2017).

26. Peterfi, L., Yusenko, M. V., Kovacs, G. & Beothe, T. FAPα and αSMA mark different subsets of fibroblasts in normal kidney and conventional renal cell carcinoma. Neoplasia 35, 100854 (2023).

27. Ho, K. H. & Patrizi, A. Assessment of common housekeeping genes as reference for gene expression studies using RT-qPCR in mouse choroid plexus. Sci Rep 11, 3278 (2021).

28. Hersom, M., Goldeman, C., Pretzer, N. & Brodin, B. Validation of reference genes for normalization of real-time quantitative PCR studies of gene expression in brain capillary endothelial cells cultured in vitro. Mol Cell Neurosci 93, 27–35 (2018).

29. Wellbrock, C., Rana, S., Paterson, H., Pickersgill, H., Brummelkamp, T. & Marais, R. Oncogenic BRAF regulates melanoma proliferation through the lineage specific factor MITF. PLoS One 3, e2734 (2008).

30. Sopel, J., Sarad, K., Kozinska, A., Mokrzyński, K., Szczygieł, D., Murzyn, A., Drzał, A., Słomiński, A., Szczygieł, M. & Elas, M. Pmel17 Deficiency Affects Melanogenesis and Promotes Tumor Vascularization. Int J Mol Sci 27, 1147 (2026).

31. Chen, Y., Yang, S., Lovisa, S., Ambrose, C. G., McAndrews, K. M., Sugimoto, H. & Kalluri, R. Type-I collagen produced by distinct fibroblast lineages reveals specific function during embryogenesis and Osteogenesis Imperfecta. Nat Commun 12, 7199 (2021).

32. Cao, J., Zheng, Z., Sun, D., Chen, X., Cheng, R., Lv, T., An, Y., Zheng, J., Song, J., Wu, L. & Yang, C. Decoder-seq enhances mRNA capture efficiency in spatial RNA sequencing. Nat Biotechnol 42, 1735–1746 (2024).

33. Zhu, J., Pang, K., Hu, B., He, R., Wang, N., Jiang, Z., Ji, P. & Zhao, F. Custom microfluidic chip design enables cost-effective three-dimensional spatiotemporal transcriptomics with a wide field of view. Nat Genet 56, 2259–2270 (2024).

34. Jerby-Arnon, L., Shah, P., Cuoco, M. S., Rodman, C., Su, M.-J., Melms, J. C., Leeson, R., Kanodia, A., Mei, S., Lin, J.-R., Wang, S., Rabasha, B., Liu, D., Zhang, G., Margolais, C., Ashenberg, O., Ott, P. A., Buchbinder, E. I., Haq, R., Hodi, F. S., Boland, G. M., Sullivan, R. J., Frederick, D. T., Miao, B., Moll, T., Flaherty, K. T., Herlyn, M., Jenkins, R. W., Thummalapalli, R., Kowalczyk, M. S., Cañadas, I., Schilling, B., Cartwright, A. N. R., Luoma, A. M., Malu, S., Hwu, P., Bernatchez, C., Forget, M.-A., Barbie, D. A., Shalek, A. K., Tirosh, I., Sorger, P. K., Wucherpfennig, K., Van Allen, E. M., Schadendorf, D., Johnson, B. E., Rotem, A., Rozenblatt-Rosen, O., Garraway, L. A., Yoon, C. H., Izar, B. & Regev, A. A Cancer Cell Program Promotes T Cell Exclusion and Resistance to Checkpoint Blockade. Cell 175, 984–997.e24 (2018).

35. Hicks, W. H., Gattie, L. C., Shami, M. E., Traylor, J. I., Davar, D., Najjar, Y. G., Richardson, T. E., McBrayer, S. K. & Abdullah, K. G. Matched three-dimensional organoids and two-dimensional cell lines of melanoma brain metastases mirror response to targeted molecular therapy. Sci Rep 14, 24843 (2024).

36. King, A. J., Arnone, M. R., Bleam, M. R., Moss, K. G., Yang, J., Fedorowicz, K. E., Smitheman, K. N., Erhardt, J. A., Hughes-Earle, A., Kane-Carson, L. S., Sinnamon, R. H., Qi, H., Rheault, T. R., Uehling, D. E. & Laquerre, S. G. Dabrafenib; preclinical characterization, increased efficacy when combined with trametinib, while BRAF/MEK tool combination reduced skin lesions. PLoS One 8, e67583 (2013).

37. Parmenter, T. J., Kleinschmidt, M., Kinross, K. M., Bond, S. T., Li, J., Kaadige, M. R., Rao, A., Sheppard, K. E., Hugo, W., Pupo, G. M., Pearson, R. B., McGee, S. L., Long, G. V., Scolyer, R. A., Rizos, H., Lo, R. S., Cullinane, C., Ayer, D. E., Ribas, A., Johnstone, R. W., Hicks, R. J. & McArthur, G. A. Response of BRAF-mutant melanoma to BRAF inhibition is mediated by a network of transcriptional regulators of glycolysis. Cancer Discov 4, 423–433 (2014).

38. Lodi, A., Woods, S. M. & Ronen, S. M. MR-detectable metabolic consequences of mitogen-activated protein kinase kinase (MEK) inhibition. NMR Biomed 27, 700–708 (2014).

39. Smith, M. P., Sanchez-Laorden, B., O’Brien, K., Brunton, H., Ferguson, J., Young, H., Dhomen, N., Flaherty, K. T., Frederick, D. T., Cooper, Z. A., Wargo, J. A., Marais, R. & Wellbrock, C. The immune microenvironment confers resistance to MAPK pathway inhibitors through macrophage-derived TNFα. Cancer Discov 4, 1214–1229 (2014).

40. Konieczkowski, D. J., Johannessen, C. M., Abudayyeh, O., Kim, J. W., Cooper, Z. A., Piris, A., Frederick, D. T., Barzily-Rokni, M., Straussman, R., Haq, R., Fisher, D. E., Mesirov, J. P., Hahn, W. C., Flaherty, K. T., Wargo, J. A., Tamayo, P. & Garraway, L. A. A melanoma cell state distinction influences sensitivity to MAPK pathway inhibitors. Cancer Discov 4, 816–827 (2014).

41. Shao, Y., Le, K., Cheng, H. & Aplin, A. E. NF-κB Regulation of c-FLIP Promotes TNFα-Mediated RAF Inhibitor Resistance in Melanoma. J Invest Dermatol 135, 1839–1848 (2015).

42. Tang, F., Lao, K. & Surani, M. A. Development and applications of single-cell transcriptome analysis. Nat Methods 8, S6–S11 (2011).

43. Kleshchevnikov, V., Shmatko, A., Dann, E., Aivazidis, A., King, H. W., Li, T., Elmentaite, R., Lomakin, A., Kedlian, V., Gayoso, A., Jain, M. S., Park, J. S., Ramona, L., Tuck, E., Arutyunyan, A., Vento-Tormo, R., Gerstung, M., James, L., Stegle, O. & Bayraktar, O. A. Cell2location maps fine-grained cell types in spatial transcriptomics. Nat Biotechnol 40, 661–671 (2022).

44. Han, Y., Wang, Y., Dong, X., Sun, D., Liu, Z., Yue, J., Wang, H., Li, T. & Wang, C. TISCH2: expanded datasets and new tools for single-cell transcriptome analyses of the tumor microenvironment. Nucleic Acids Res 51, D1425–D1431 (2023).

