## Supplementary material for "Live Spatial Transcriptomics Enables Longitudinal Tissue Mapping": SI figure

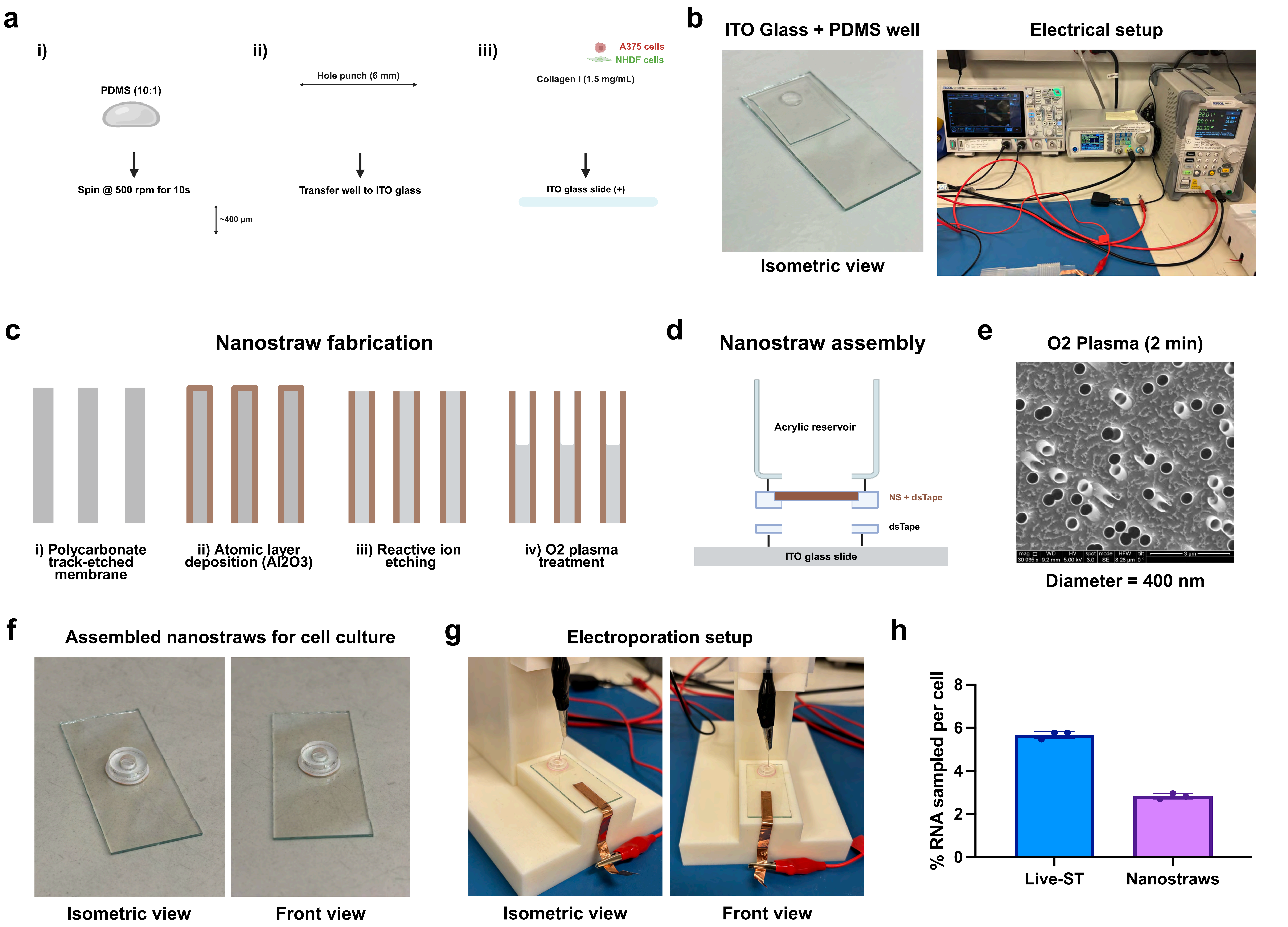


**Supplementary Figure 1: 3D hydrogel intracellular sampling.** (**a**) Fabrication of the three-dimensional (3D) co-culture model: (i) polydimethylsiloxane (PDMS; 10:1 base: curing agent) was spin-coated onto a glass slide at 500 rpm for 10 s to a thickness of ~400 µm; (ii) a 6 mm well was punched and transferred to indium tin oxide (ITO) glass; (iii) human melanoma cells (A375) and normal human dermal fibroblasts (NHDF) in collagen I (1.5 mg/mL) were seeded into the well and electroporated with a top ITO glass slide. Created with BioRender.com. (**b**) Photographs of the ITO glass with PDMS well (left) and the electrical setup (right). (**c**) Nanostraw fabrication: (i) polycarbonate track-etched membrane, (ii) atomic layer deposition of $Al_2O_3$, (iii) reactive ion etching, and (iv) O2 plasma treatment. (**d**) Nanostraw assembly with an acrylic reservoir and double-sided tape on ITO glass. (**e**) Scanning electron micrograph of nanostraws after 2 min of O2 plasma treatment (diameter, 400 nm). (**f**) Assembled nanostraws for cell culture. (**g**) Nanostraw electroporation setup. (**h**) RNA sampled per cell with Live Spatial Transcriptomics (Live-ST) and with nanostraws. Data in (**h**) are shown as mean ± SD (*n* = 3 technical replicates). Scale bar: 3 µm (**e**).

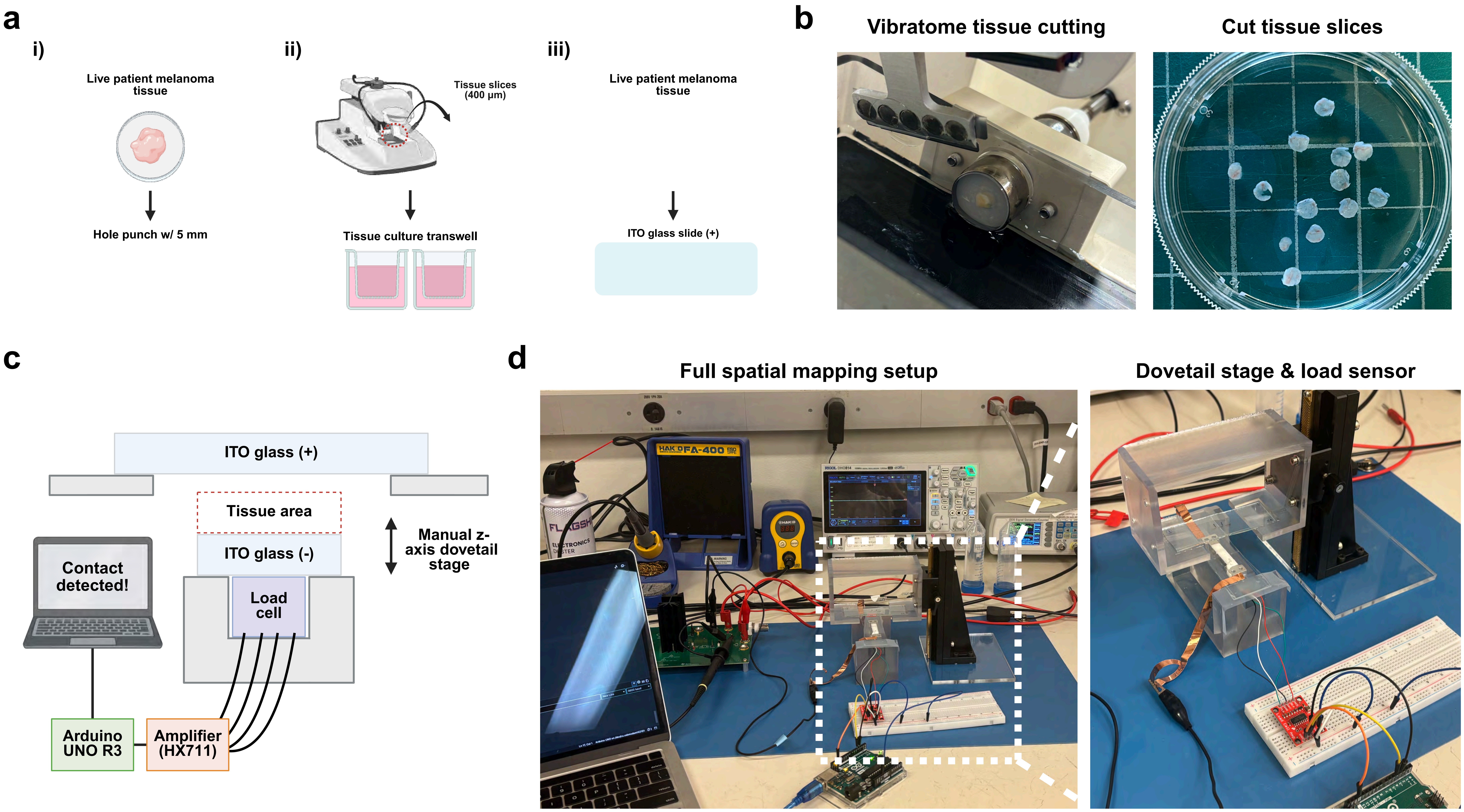


**Supplementary Figure 2: Live tissue preparation and spatial mapping setup.** (**a**) Tissue preparation: (i) patient-derived melanoma tissue was cored with a 5 mm biopsy punch; (ii) cores were sectioned into 400 µm slices with a vibratome and cultured on Transwell inserts; (iii) slices were placed between ITO glass slides for Live-ST sampling. Created with BioRender.com. (**b**) Vibratome sectioning (left) and cut tissue slices (right). (**c**) Schematic of the spatial mapping setup. A load cell beneath the tissue-supporting ITO slide, read out through an HX711 amplifier and an Arduino UNO R3, detects contact between the capture slide and the tissue as the slide is lowered with a manual z-axis dovetail stage. (**d**) Photographs of the full spatial mapping setup (left) and the dovetail stage with load cell (right).

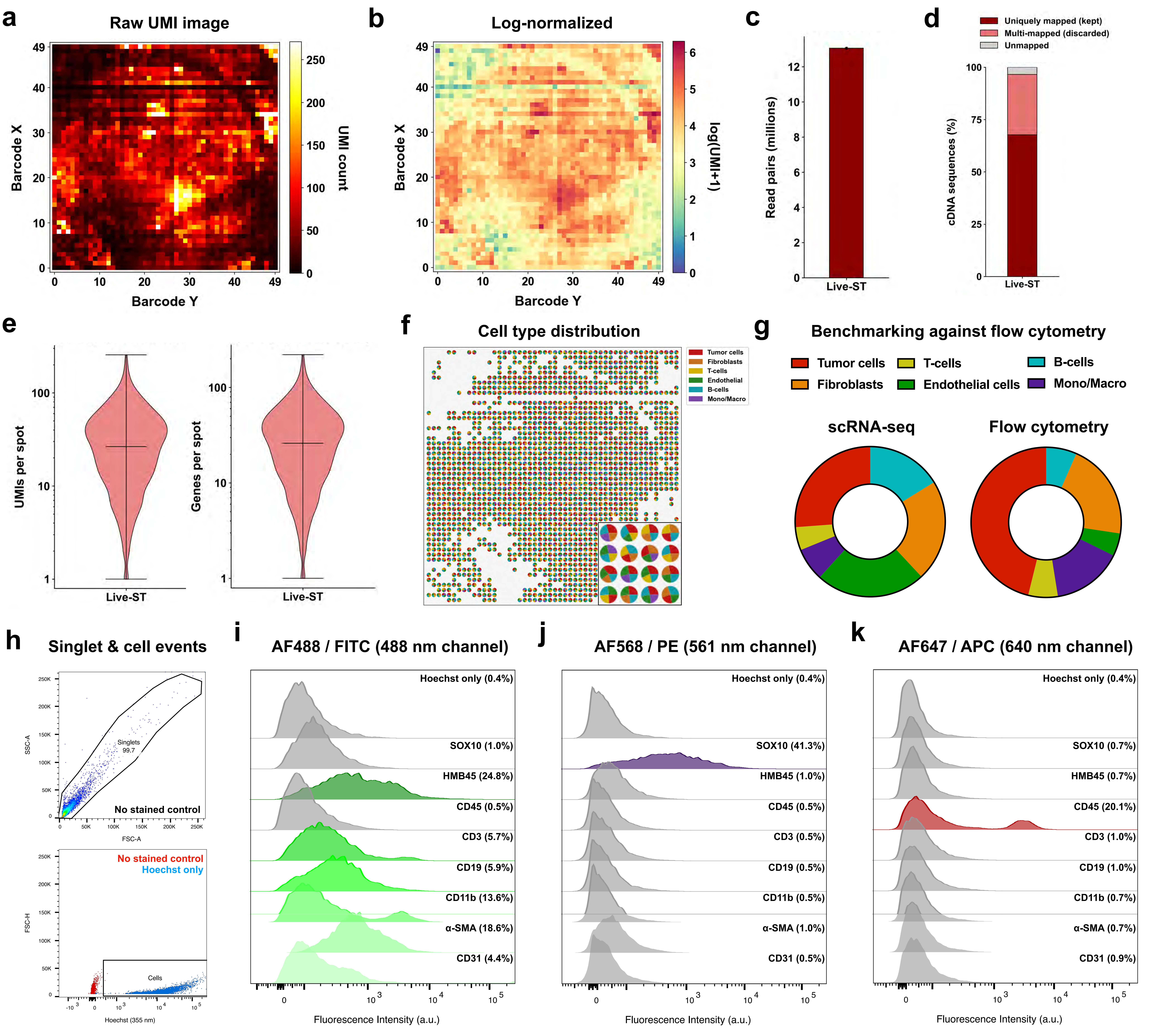


**Supplementary Figure 3: Live-ST and cell type mapping validation.** (**a**,**b**) Raw unique molecular identifier (UMI) counts (**a**) and log-normalized expression, log(UMI + 1) (**b**), per spot for the melanoma slice in **Figure 3i** (Patient 2). (**c**) Number of read pairs. (**d**) Fraction of cDNA sequences that were uniquely mapped, multi-mapped (discarded), or unmapped. (**e**) UMIs (left) and genes (right) per spot, counted from gene-assigned molecules, across all spots with data. Spots with at least 25 uniquely mapped UMIs were retained for analysis. (**f**) The four most abundant cell types in each mapped spot (rescaled to sum to 100%), shown as pie charts; the inset shows an enlarged region. (**g**) Cell type proportions estimated from Live-ST of Patient 2 spots mapped with the single-cell RNA sequencing (scRNA-seq) reference (left, "scRNA-seq") and measured by flow cytometry of Patient 3 (right). Flow cytometry proportions were scaled to sum to 100% across the six cell types. (**h**) Gating of cells (forward scatter area (FSC-A) versus side scatter area (SSC-A)) to exclude debris, followed by Hoechst-positive cell events, using unstained and Hoechst-only controls. (**i**–**k**) Fluorescence histograms of tumor (SOX10, HMB45), immune (CD45, CD3, CD19, CD11b), fibroblast (α-SMA), and endothelial (CD31) markers in the Alexa Fluor (AF) 488/fluorescein isothiocyanate (FITC) (**i**), AF568/phycoerythrin (PE) (**j**), and AF647/allophycocyanin (APC) (**k**) channels. Percentages indicate marker-positive events.

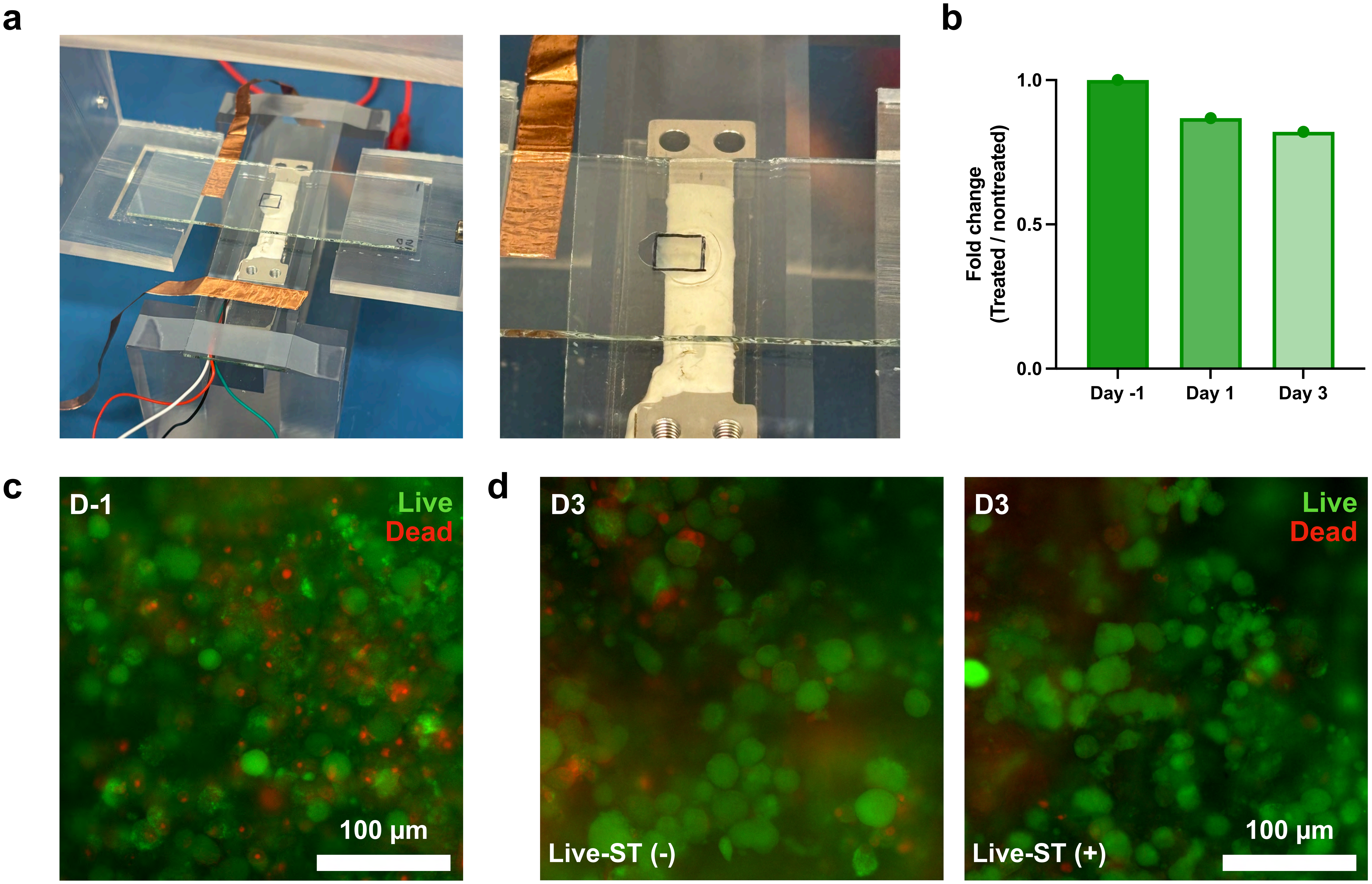


**Supplementary Figure 4: Setup and cell viability assessment for the longitudinal study.** (**a**) Photographs of the Live-ST sampling setup used to repeatedly sample tissue slices. (**b**) Metabolic activity of drug-treated relative to nontreated viability-control slices one day before treatment (D-1) and on days 1 and 3 after treatment (D1 and D3), measured by MTS assay and shown as fold change. (**c**) Live/Dead staining of a tissue slice at D-1. (**d**) Live/Dead staining at D3 of slices cultured without (Live-ST (−)) or with (Live-ST (+)) repeated sampling. Live cells are shown in green and dead cells in red. All tissue is from Patient 4. Scale bars: 100 µm (**c**,**d**).

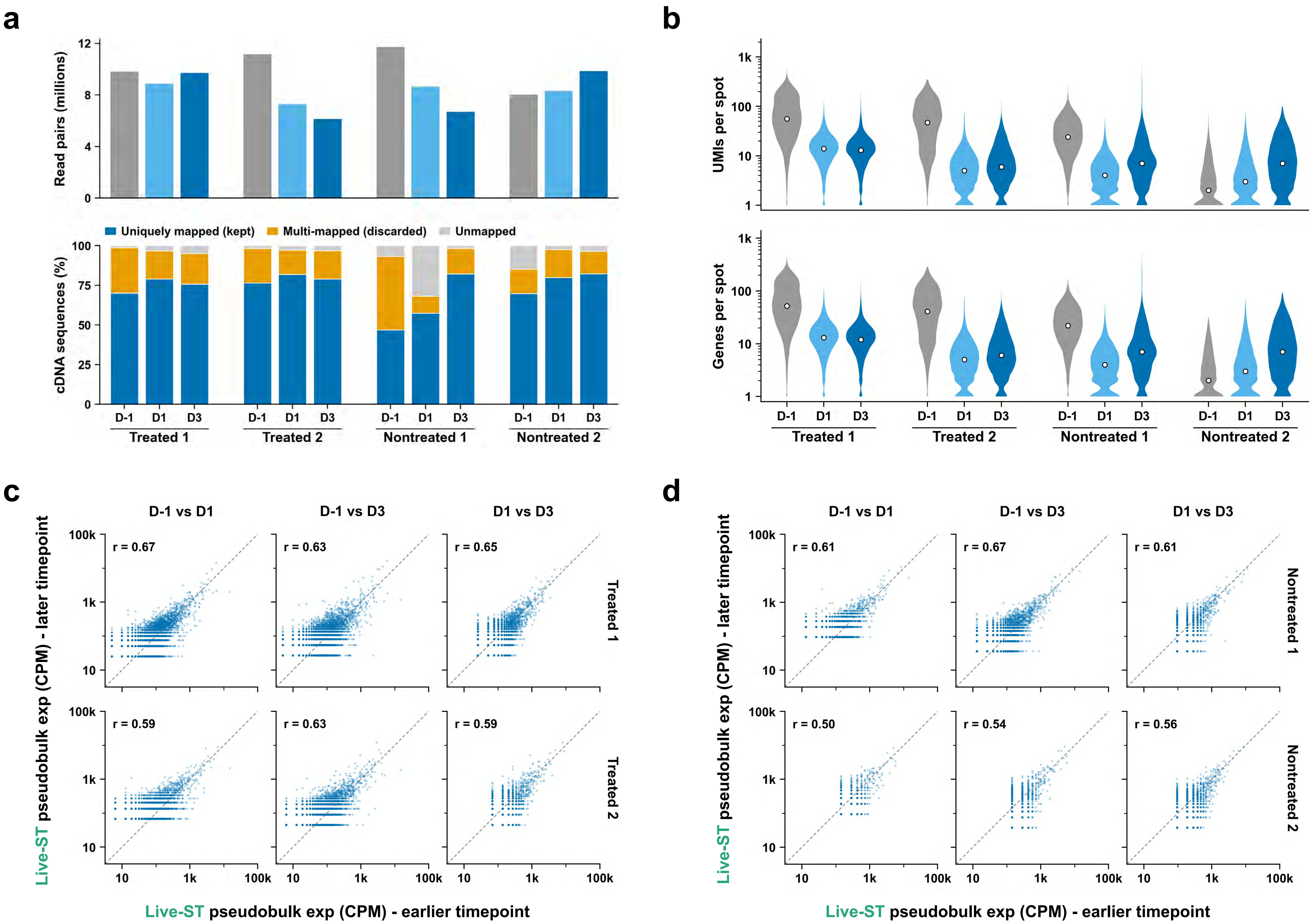


**Supplementary Figure 5: Quality control for longitudinal profiling of drug responses.** (**a**) Number of read pairs (top) and fraction of cDNA sequences that were uniquely mapped, multi-mapped (discarded), or unmapped (bottom) for the 12 libraries from Treated 1 and 2 and Nontreated 1 and 2 at D-1, D1, and D3. (**b**) UMIs (top) and genes (bottom) per spot for each library, counted from gene-assigned molecules over all spots with data; white dots indicate medians. (**c**,**d**) Pseudobulk expression (counts per million, CPM; UMIs summed over tissue spots) between time points within treated (**c**) and nontreated (**d**) tissues, excluding ribosomal protein and mitochondrial genes. r, Pearson's correlation coefficient of log10 CPM over genes detected in both samples. The dashed line indicates the identity line. All tissue is from Patient 4.
